# ProlateNet: Sampling-rate transfer of convolutional networks with a weights-only certificate for physiological signals

**DOI:** 10.64898/2026.11.17.767313

**Authors:** Han Cui, Yuge Zhu, Bing Li, Shaojia Huang, Tao Xu

**Affiliations:** Department of Biomedical Engineering, College of Engineering, Shantou University, 243 Daxue Road, Shantou 515063, China; Department of Electronic Engineering, College of Engineering, Shantou University, 243 Daxue Road, Shantou 515063, China; School of Intelligent Manufacturing and Aeronautics, Zhuhai College of Science and Technology, 8 Anji East Road, Jinwan District, Zhuhai 519041, China

**Keywords:** Sampling-rate transfer, Continuous-kernel convolution, Prolate spheroidal wave functions, Aliasing, Physiological signals, Electroencephalography, Electrocardiography

## Abstract

A central goal of physiological signal analysis is to train a model once and deploy it across recording systems. However, these systems sample EEG and ECG at different rates, and a network trained at one rate loses accuracy at the others. Continuous-kernel networks were built to solve this problem, but published ones still fail at unseen rates, and no existing method can certify a trained model before deployment. In this study, we propose ProlateNet, which builds kernels from prolate spheroidal wave functions to suppress aliasing across sampling rates and certifies a trained model from its weights alone. Across fourteen first-layer designs trained at one rate on three public EEG and ECG datasets, the published continuous-kernel layers FlexConv and sampling-frequency-independent (SFI) convolution show mean worst drops of 0.18 and 0.35 macro-F1 on SEED. ProlateNet is the only design that is stable and accurate on all three datasets with an error bound fixed before training, and its mean worst drop is at most 0.003. Without targetrate recordings, its certificate ranks the continuous-kernel designs on SEED by measured transfer with Spearman ρ = 0.92, against 0.68 for energy above the training Nyquist frequency. Overall, ProlateNet shows that controlling aliasing enables convolutional networks to transfer across sampling rates and provides principled guidance for building physiological signal models that are trained once and deployed across recording systems.

## 1. Introduction

Deep networks now decode brain states, sleep stages and cardiac activity from physiological recordings [1–4], and a central goal of the field is to train such a model once and deploy it across recording systems. However, these systems sample at different rates [4–7], and a network trained at one rate loses accuracy at the others. The cause lies in its convolutional filters: their taps are fixed on the training grid, so a change in sampling rate stretches or compresses them in physical time and shifts their frequency response [1,2]. Existing pipelines therefore resample recordings to a common rate [3,8,9] or fine-tune models on new recordings [10], at the cost of extra filter delay or new labelled data. Continuous-kernel methods avoid this cost by defining each kernel in continuous time and sampling it at the deployment rate, which preserves its physical time scale and reuses the learned parameters [11–13].

However, preserving a kernel’s physical time scale does not by itself ensure consistent filtering across sampling rates. When a continuous kernel is sampled, its high-frequency energy can fold back into the signal band, producing aliasing [13]. Because the frequencies that fold back depend on the sampling rate, the same continuous kernel produces different in-band responses at different rates. Flex-Conv controls kernel frequencies up to the Nyquist frequency of the training rate [13], which leaves lower deployment rates unprotected. Reliable sampling-rate transfer therefore calls for a quantitative link between kernel spectral properties and filtering error across all deployment rates.

Before using a trained model at a new sampling rate, we also need to assess how much its filtering response will change. For audio source separation, Imamura et al. assess sampling-frequency independence by measuring how network components respond to resampling [14], which requires inputs at every tested rate. To assess the first layer without target-rate inputs, we seek an error bound that can be calculated directly from the trained kernel weights. We can then compare this bound with an acceptable error level before testing the kernel at a new sampling rate.

Beyond assessing trained kernels, we also need to choose their bandwidth and number of basis functions before training. A narrow bandwidth restricts the temporal patterns a kernel can represent, while a wider bandwidth can admit frequencies that produce aliasing [13]. Likewise, the number of well-concentrated basis functions is constrained by the kernel’s duration and bandwidth, as classical time– bandwidth theory shows [15]. We therefore need a design rule that selects band-width and basis size to satisfy a specified error tolerance across deployment rates.

To address these challenges, we propose ProlateNet, a new framework for transferring convolutional networks across sampling rates without retraining. We derive a bound on offset-averaged first-layer aliasing error that holds uniformly across admissible sampling rates. Guided by this bound, ProlateNet represents continuous kernels using prolate spheroidal wave functions (PSWFs) and samples them at exact time lags [15,16]. The same bound determines kernel bandwidth and basis size and can be evaluated directly from the trained weights before deployment. We use the normalized out-of-band kernel energy as a transfer certificate for first-layer error assessment.

Our contributions are as follows:

1. We propose ProlateNet, a framework that enables convolutional networks trained at one sampling rate to operate at unseen rates without retraining or input resampling.
2. We derive a uniform upper bound on the offset-averaged first-layer aliasing error. This bound determines kernel bandwidth and basis size and yields a transfer certificate calculated from trained weights before deployment.
3. We compare fourteen approaches to first-layer design on three public EEG and ECG datasets and demonstrate stable transfer with competitive classification performance. Further experiments show how the kernel basis and theoretically chosen bandwidth and basis size affect transfer.

## 2. Related work

This section reviews three areas relevant to ProlateNet: convolutional kernel design, sampling-rate transfer and its evaluation, and aliasing and spectral concentration.

### 2.1. Convolutional kernel design

Existing studies have introduced structure into convolutional kernels through parametric filters and basis expansions. For example, SincNet learns the lower and upper cutoff frequencies of band-pass filters [17], and Borra et al. applied this parameterization to EEG decoding of executed and imagined movements [18].

LEAF [19] and adaptive Gabor convolutional networks [20] learn Gabor filters for audio and image analysis, respectively. Beyond learning filter parameters directly, another approach constructs convolutional kernels by learning combinations of basis functions. For example, DCFNet expresses kernels as weighted sums of fixed basis functions, including Fourier–Bessel functions, and learns the expansion coefficients [21]. These approaches show how the choice of filter family or basis can control kernel structure while reducing the number of trainable parameters.

Building on these structured designs, continuous-kernel methods further consider how kernels can accommodate different sampling grids. CKConv represents kernels as continuous functions generated by a small neural network, allowing their values to be evaluated at different sampling locations [12]. FlexConv further introduces learnable kernel sizes and a parameterization that allows analytic control of kernel frequencies to reduce aliasing [13]. For audio source separation, Saito et al. construct sampling-frequency-independent (SFI) layers by discretizing analog filters at the deployment sampling rate [11,22]. Imamura et al. extend this approach by representing the analog filters with neural networks [23]. These continuous formulations allow kernel values to be evaluated on different sampling grids while retaining the learned parameters.

### 2.2. Sampling-rate transfer and its evaluation

Sampling-rate transfer is part of the broader problem of applying learned models on different discretization grids. Neural operators learn mappings between function spaces and support evaluation on different grids [24,25], while representation-equivalent neural operators formalize conditions for alias-free operator learning [26]. For sequential data, structured state-space models such as S4 adjust their discretization step to accommodate changes in sampling rate [27]. Related timeseries studies address differences in observation times by aligning variables recorded at multiple rates [28] or reconstructing temporal patterns in irregularly sampled data [29]. These studies connect continuous representations with learning from observations acquired on different grids.

In physiological signal analysis, combining recordings from multiple sources also requires accommodating differences in acquisition. U-Sleep, BIOT and LaBraM resample recordings to a common rate before model processing [3,8,9]. Other studies learn representations for cross-device EEG emotion recognition [30] or align features across subjects and sessions [31]. For evaluating sampling-rate transfer itself, Imamura et al. introduce local equivariance error metrics that measure how network components respond to resampling in audio source separation [14]. Such response-based metrics complement task-level measures by directly examining changes in network outputs across sampling rates.

### 2.3. Aliasing and spectral concentration

Research on aliasing in convolutional networks has mainly examined the effects of downsampling on learned representations. Azulay and Weiss relate sensitivity to small image transformations to sampling effects [32], and Zhang introduces low-pass filtering before downsampling to improve shift invariance [33]. Karras et al. treat feature maps as continuous signals when designing alias-free generators [34], while Xu et al. use Haar wavelet downsampling to preserve information during resolution reduction [35]. Together, these studies demonstrate the importance of controlling frequency content when discrete representations are sampled or downsampled.

Classical time–bandwidth theory provides a mathematical basis for controlling spectral concentration. For functions confined to a fixed time window, prolate spheroidal wave functions provide basis functions ordered by their energy concentration within a prescribed frequency band [15,16]. Landau and Widom characterize the associated eigenvalue distribution, which determines the number of well-concentrated functions [36]. Complementing these concentration results, Blu and Unser analyse offset-averaged approximation error in the Fourier domain [37]. PSWFs have also been used as activation functions in implicit neural representations [38], connecting this classical theory with neural function approximation.

Although continuous kernels and frequency control improve adaptability to different sampling grids, quantifying filtering error before deployment requires a direct link between kernel spectra and sampling-rate transfer error. To this end, we design ProlateNet to combine continuous-kernel design with a uniform bound on offset-averaged first-layer aliasing error across admissible sampling rates. By linking kernel design to an explicit error bound, ProlateNet provides a theoretical basis and a practical approach for deploying one trained model across sampling rates without retraining.

## 3. Problem formulation

### 3.1. Setting and objective

We consider physiological signal classification when the sampling rate changes between training and deployment. A network is trained once at rate *f*_0_ and subsequently applied at a target rate *f*. We denote the lowest rate covered by training and deployment as *f*_min_. Our objective is to reuse the trained network across these rates without retraining or resampling the input recordings, while maintaining consistent filtering behavior in the first temporal layer.

To describe the same filter across sampling rates, we represent each temporal kernel as a continuous function *k*(*t*) supported on a window of duration *D*. We assume that the first-layer input is band-limited to [−*B, B*], with *f*_min_ > 2*B*. At rate *f*, the continuous kernel is evaluated at physical time lags *n*/*f* to obtain discrete convolution coefficients *h*_*f*_ [*n*], with scaling that accounts for the sampling interval. The same kernel parameters are therefore retained across rates, while the discrete coefficients are generated on the corresponding sampling grid.

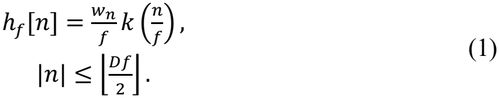

The endpoint weight is *w*_*n*_ = 1/2 when |*n*|/*f* = *D*/2 and *w*_*n*_ = 1 otherwise; Section 4.4 details this realization.

However, evaluating a continuous kernel on different grids does not by itself ensure consistent filtering responses. We describe the response at rate *f* by

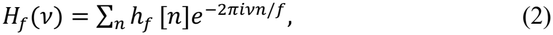

where *v* denotes frequency in hertz. We compare this response with the continuous kernel spectrum *K*(*v*). We quantify the discrepancy by integrating the squared response error over the signal band and averaging over shifts of the kernel relative to the sampling grid. Our theoretical objective is to bound this offset-averaged first-layer error uniformly for all *f* ≥ *f*_min_, using quantities computable from the kernel parameters.

### 3.2. Overall framework

To achieve this objective, ProlateNet combines a continuous first-layer kernel with a theoretical error bound, as illustrated in Fig. 1. The error analysis identifies the kernel frequencies that contribute to aliasing across deployment rates. Guided by this analysis, we construct kernels from prolate spheroidal wave functions and determine their bandwidth and number of basis functions. We then evaluate these kernels at the exact time lags of each sampling rate, connecting the continuous design to its discrete implementation.

**Fig. 1.**
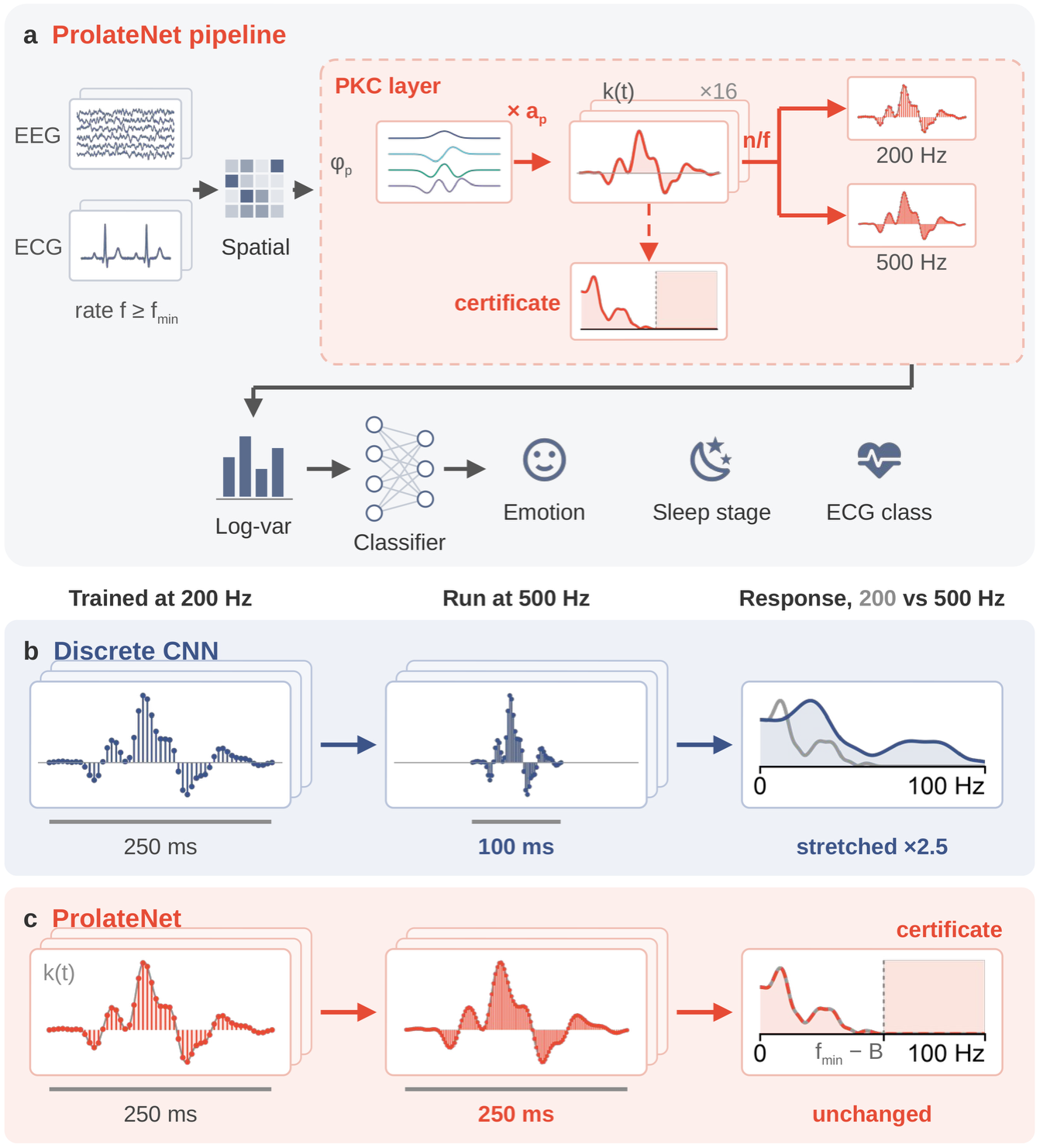
ProlateNet keeps its first-layer kernels across sampling rates, whereas a discrete CNN rescales them. (a) A recording at any rate *f* ≥ *f*_min_ passes through a spatial projection, the prolate kernel convolution (PKC) layer, mean and log-variance statistics over 1-s segments and a classifier, which outputs emotion, sleep-stage or ECG labels. The PKC layer weights prolate spheroidal wave functionsφ_*p*_ by coefficients *a*_*p*_ into a continuous kernel *k*(*t*) and samples it at the exact lags *n*/*f* of each rate. Solid arrows mark the forward pass, and the dashed arrow marks the transfer certificate, which is computed from the same coefficients before deployment. (b, c) Both networks are trained at 200 Hz and run unchanged at 500 Hz. (b) A discrete CNN stores 51 taps. At 500 Hz the same taps span 100 ms instead of 250 ms, so the frequency response stretches by a factor of 2.5 and the filter moves away from the band it learned. (c) ProlateNet keeps the duration and frequency response of *k*(*t*) at both rates. The shaded band above *f*_min_ − *B* is the kernel energy that the certificate measures. Frequency responses are plotted up to 100 Hz, the Nyquist frequency at 200 Hz. Stacked cards stand for the 16 kernels of the first layer. Kernels are drawn with one fixed set of illustrative weights, and the EEG and ECG traces are schematic.

During training, a spatial projection first combines the input channels, and the prolate kernel convolution (PKC) layer extracts temporal features from the resulting signals. The network then computes the mean and log-variance of each output map over fixed-duration segments and averages these statistics across segments. A two-layer perceptron with layer normalization and Gaussian error linear unit (GELU) activation uses the resulting features to predict the physiological signal labels. The statistics are computed over 1-s segments. The spatial projection produces two maps per filter, and each temporal kernel filters its associated pair of maps. The kernel expansion coefficients, spatial projection and classifier are learned jointly from recordings at *f*_0_.

At deployment, we generate the discrete kernel coefficients for the target rate *f* from the trained continuous kernels, while retaining the learned expansion coefficients and all other network weights. The segment statistics provide features of the same dimension for the classifier across sampling rates. Before processing target-rate recordings, we also compute the first-layer aliasing-error bound directly from the trained kernel weights. Section 4 develops the error analysis, kernel design and deployment procedure in detail.

## 4. ProlateNet

### 4.1. Sampling-rate transfer error

To design kernels for sampling-rate transfer, we first identify which spectral components control the filtering error. For a kernel of bounded variation, the Poisson summation formula expresses its discrete response as a sum of shifted copies of the continuous kernel spectrum, with midpoint values used at the window endpoints [39]. The unshifted spectrum gives the intended continuous response, while the remaining copies contribute aliasing.

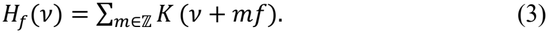

For an input confined to the signal band, only kernel frequencies beyond the sampling rate minus the signal bandwidth can fold into that band. Across deployment rates, the lowest rate therefore determines a common frequency boundary, given by *f*_min_ − *B*. Controlling kernel energy beyond this boundary provides a shared design target for the entire deployment range.

However, the alias components add in amplitude, so their combined effect also depends on the kernel’s position relative to the sampling grid. We account for this dependence by comparing the discrete and continuous responses within the signal band and averaging the squared discrepancy over one sampling interval. This defines the offset-averaged first-layer response error introduced in Section 3.

Let *θ* ∈ [0,1) shift the physical lags to (*n* + *θ*)/*f*, and let *H*_*f,θ*_ denote the response referred to these physical lags. We define

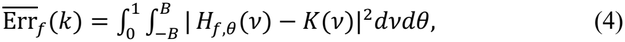

Write *ℰ*_Ω_ (*k*) = *∫*_Ω_ | *K*(*ξ*)|^2^*dξ* and *A*_*f*_ = ⋃_*m≠ 0*_(*mf* + [−*B, B*]).

#### Proposition 1

(Appendix A.2). For every square-integrable kernel supported on [−*D*/2, *D*/2] and every *f* ≥ *f*_min_ > 2*B*,

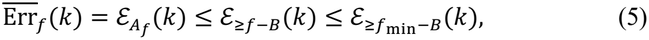

Averaging over the sampling offset removes cross terms between different alias orders, allowing their energies to add. Classical Fourier error analysis provides this energy identity [37], and the inclusion of the alias bands within the common frequency region gives the uniform bound. Consequently, errors at different deployment rates can be controlled through the same kernel spectral quantity. The proof is provided in Appendix A.2.

For a nonzero kernel, we normalize the spectral bound by its energy,

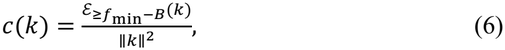

For a linear kernel space *V*, write *c*(*V*) = sup_0≠*k*∈*V*_*c*(*k*). This measures the worst-case out-of-band energy fraction over that space.

Proposition 1 gives a direct design criterion: concentrate kernel energy within the band bounded by *f*_min_ − *B*. We use this criterion to select a kernel representation whose spectral concentration can be controlled during design and calculated after training. The following subsection introduces the prolate spheroidal wave function parameterization.

### 4.2 Prolate kernel parameterization

The error bound motivates kernels that concentrate their energy within the prescribed frequency band. At the same time, each kernel must operate within a finite time window and retain sufficient flexibility for learning temporal features. We therefore use prolate spheroidal wave functions, which maximize spectral concentration under a fixed time-window constraint [15,16].

For a prescribed duration and bandwidth, we compute the basis functions before training and order them by their in-band energy concentration. Each kernel is represented as a weighted sum of the first *P* basis functions, with a fixed gain that normalizes its initial discrete coefficients at the training rate. Training learns the expansion coefficients, while the basis functions and gains remain fixed across sampling rates.

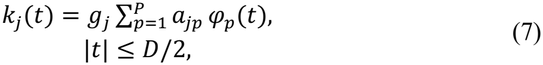

Here *a*_*jp*_ are learnable expansion coefficients and *g*_*j*_ is the fixed gain of filter *j*; the functions *φ*_*p*_ form the precomputed basis.

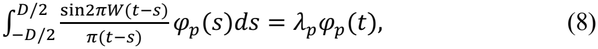

The functions *φ*_*p*_ are orthonormal on [−*D*/2, *D*/2], with concentration eigenvalues 1 > *λ*_5_ > *λ*_)_ > … > 0. We compute them once per (*D, W*) by Nyström discretization on 512 Gauss–Legendre nodes.

This representation connects the learned coefficients directly to the error bound. Because the basis functions are orthonormal and diagonalize the spectral concentration operator, the out-of-band kernel energy is a weighted sum of squared expansion coefficients. The weights are determined by the precomputed concentration eigenvalues. We can therefore calculate the spectral quantity identified in Section 4.1 directly from the trained coefficients.

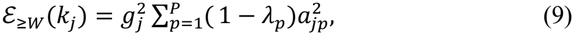

### 4.3. Bandwidth and basis size

The remaining design choices are the bandwidth *W* and the number of basis functions *P*. A narrower band restricts the temporal patterns available to the kernel, while a wider band admits more energy into frequencies that can alias. Guided by Proposition 1, we set *W* = *f*_min_ − *B*, linking the kernel bandwidth to the signal band and the intended deployment range.

This choice also has a minimax interpretation. Among all linear kernel spaces of a given dimension and duration, the leading PSWFs at this bandwidth minimize the worst-case fraction of energy beyond the common frequency boundary. This follows from the Courant–Fischer principle applied to the spectral concentration operator [40,41]. Consequently, the selected bandwidth supports the largest basis size under a prescribed tolerance on that energy fraction.

#### Proposition 2

(Appendix A.3). (i) For every *P*-dimensional space *V*of kernels supported on [−*D*/2, *D*/2], the worst-case certificate satisfies

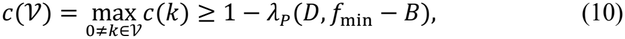

with equality for the PKC space with *W* = *f*_min_ − *B*. (ii) Consequently, at every tolerance *ε*, the band *W* = *f*_min_ − *B* admits at least as many basis functions with *c*(*V*) ≤ *ε* as any other band.

We determine *P* from the concentration eigenvalues. These eigenvalues remain close to one for approximately the time–bandwidth dimension and then decrease, limiting the number of well-concentrated basis functions [15,36]. Retaining only functions whose eigenvalues exceed the chosen concentration threshold bounds the out-of-band energy fraction of every kernel in the resulting space. The same worst-case spectral criterion applies to other linear kernel representations.

#### Proposition 3

(Lemma A.1, Appendix A.4). Let the deployment rates fill an interval from *f*_min_ to at least 2*f*_min_ − 2*B*, which is 110 Hz in our setting. Then every linear space of kernels supported on [−*D*/2, *D*/2] whose dimension exceeds *N*_1 − *ε*_ (*D, f*_min_ − *B*), the number of eigenvalues of (8) at *W* = *f*_min_ − *B* that are at least 1 − *ε*, contains a kernel that places more than a fraction *ε* of its energy in frequencies that alias at some deployment rate. The PKC space with *W* = *f*_min_ − *B* and *P* = *N*_1 − *ε*_ attains the limit.

In our experiments, a kernel duration of 250 ms, a signal bandwidth of 45 Hz and a minimum rate of 100 Hz give a kernel bandwidth of 55 Hz. Requiring at least 99% in-band concentration yields *P* = 25. With sixteen kernels, the layer therefore learns 400 expansion coefficients, independently of the deployment sampling rate.

### 4.4. Exact-lag realization

After selecting the continuous kernel space, we construct the discrete convolution at the target rate. Each kernel is evaluated at the exact physical time lags *n*/*f*, and its values are scaled by the sampling interval 1/*f*. The kernel duration therefore remains fixed in physical time as the sampling rate changes.

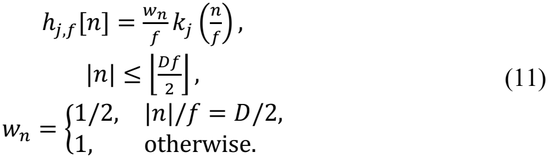

When a sampling point coincides with a window endpoint, we assign it half weight to obtain the midpoint value required by the periodization identity. This avoids an additional endpoint contribution to the discrete response. Evaluating the kernel at exact lags also preserves the intended time scale when the window edges ±D/2 do not fall on the sampling grid.

For each target rate, the basis values and sampling weights form a fixed matrix that maps the learned coefficients to discrete taps. These taps are generated once before deployment and used in a standard depthwise convolution with zero padding. The expansion coefficients, spatial projection and classifier retain their trained values.

### 4.5. Error assessment from trained weights

The same parameterization allows error assessment before processing recordings at a new sampling rate. For each trained kernel, we calculate the fraction of its energy beyond *f*_min_ − *B*. At the selected bandwidth, this fraction follows directly from the squared expansion coefficients and the precomputed concentration eigenvalues. We refer to this normalized spectral quantity as the transfer certificate.

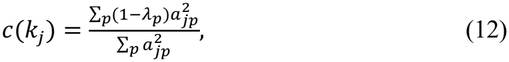

Multiplying the certificate by the kernel energy gives the uniform upper bound on its offset-averaged response error. To summarize a layer containing multiple filters, we calculate the total out-of-band energy divided by the total kernel energy. This produces an energy-weighted layer certificate using only the trained weights and fixed basis information.

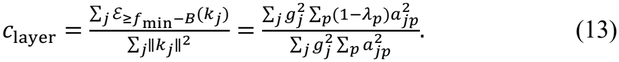

The certificate connects kernel design with assessment after training: the basis and eigenvalue threshold control the entire kernel space, while the learned coefficients determine the value for each trained model. For the implemented fixed sampling grids, the supplementary analysis additionally provides bounds with constants determined by the finite-dimensional kernel space.

### 4.6. Deployment procedure and computational cost

Deployment requires the trained expansion coefficients, fixed gains and precomputed basis functions. We first calculate the layer certificate and then generate the discrete taps at the target rate. The resulting network processes the target-rate recordings using the new taps and the unchanged remaining weights, without retraining or input resampling.

For direct convolution, both the number of input samples and the kernel length increase with the sampling rate, giving quadratic growth in first-layer computation. We therefore provide an optional strided implementation that evaluates the output at selected sampling positions. The retained outputs are identical to the corresponding outputs of the full-rate convolution.

We select an integer stride that retains an output rate above twice the signal band-width, using a practical margin of 120 Hz in our experiments. This reduces the growth in convolutional cost to approximately linear in the input sampling rate. The classifier continues to receive segment means and log-variances, and the effects of output subsampling on these features and classification performance are evaluated experimentally.

We choose the largest integer divisor *s* of *f* satisfying *f*/*s* ≥ 120 Hz. Algorithm 1 summarizes the deployment procedure.

#### Algorithm 1

Deployment of a trained PKC layer at a new rate *f*

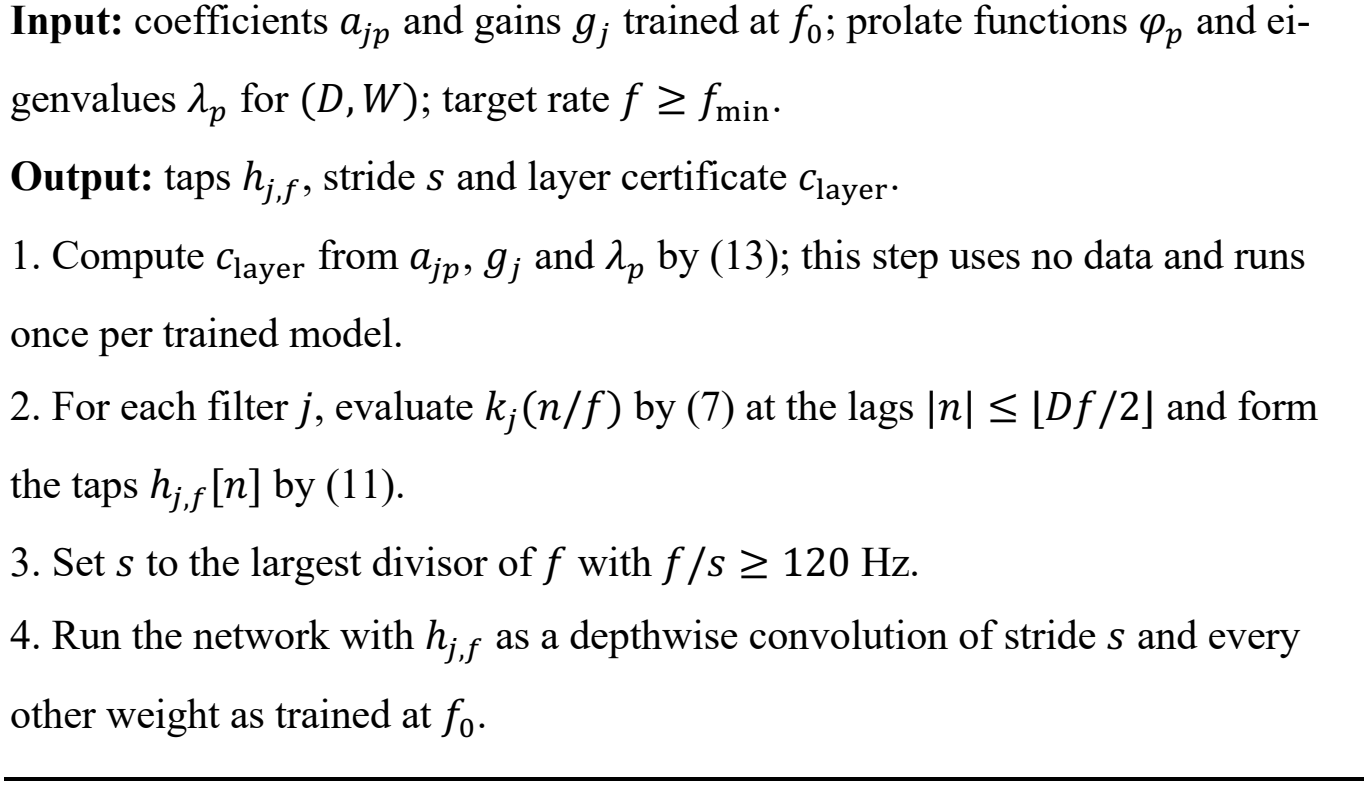

## 5. Experiments

### 5.1. Datasets, protocol and metrics

We evaluate ProlateNet on three public physiological signal datasets: SEED for EEG emotion recognition [7], Sleep-EDF-20 for sleep staging [5,6], and PTB-XL for ECG classification [4]. These datasets cover different prediction tasks, channel configurations and recording rates. Table 1 summarizes their characteristics and evaluation protocols. In the main comparison, each model is trained at a single sampling rate and evaluated at unseen target rates without retraining or finetuning.

**Table 1.**
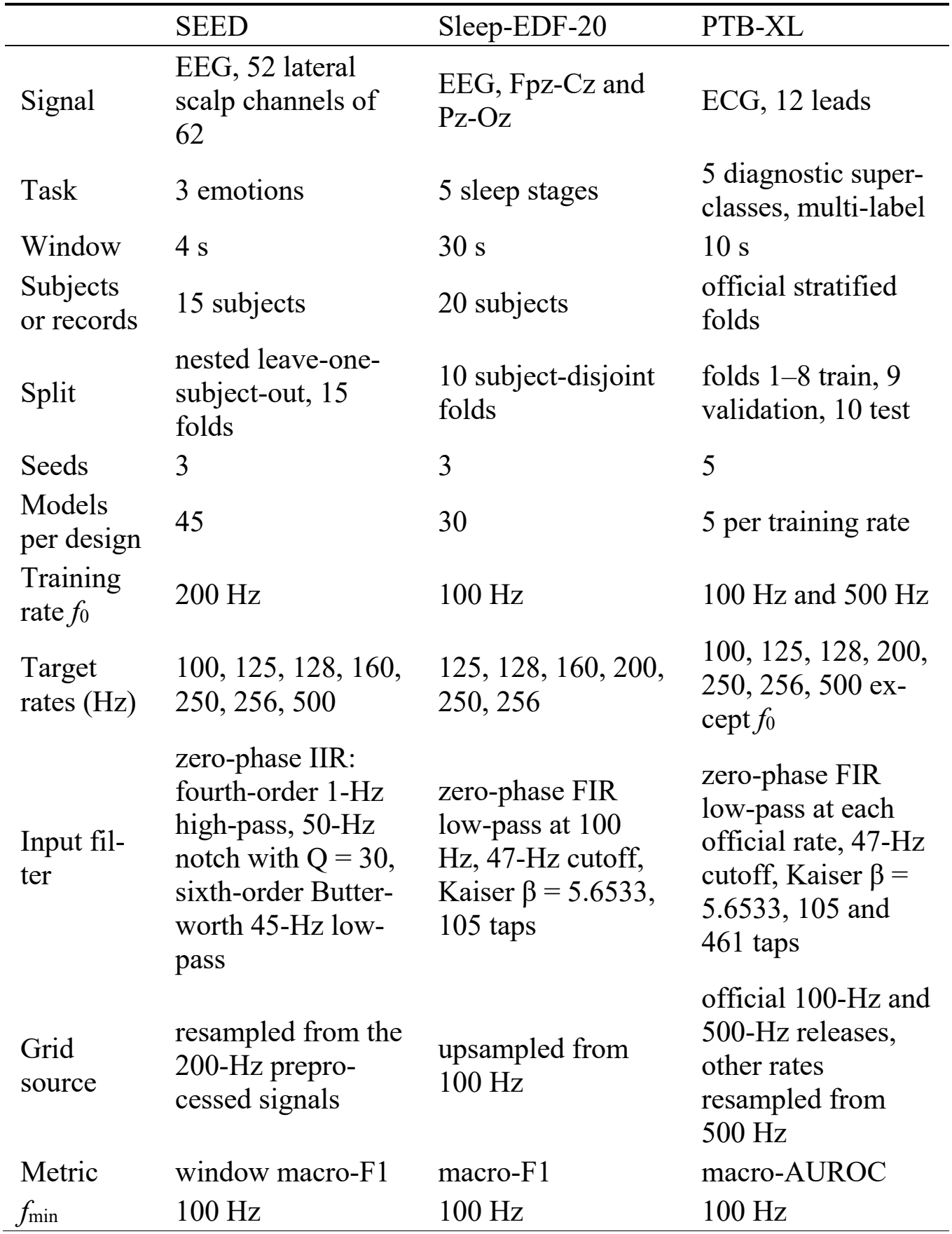
Datasets and protocol.

For SEED, we use nested leave-one-subject-out evaluation, with the held-out participant excluded from training and validation. Sleep-EDF-20 uses ten subject-disjoint folds, while PTB-XL follows its official split, using folds 1–8 for training, fold 9 for validation and fold 10 for testing. We repeat training with multiple random seeds and retain the same data partitions across sampling rates.

Before constructing the sampling grids, we restrict the analysis band to 45 Hz using the dataset-specific filters listed in Table 1. To generate recordings at additional rates, we apply polyphase FIR resampling to complete recordings before extracting the evaluation windows. This preserves the same physical windows across rates. During inference, each model directly processes the windows at their target sampling rate.

SEED models are trained at 200 Hz, and Sleep-EDF models at 100 Hz. For PTBXL, we train separate models at 100 and 500 Hz to examine transfer from both lower and higher training rates. Each trained model is evaluated at the remaining rates specified in Table 1. The lowest rate across all evaluation settings is 100 Hz, giving a common spectral threshold of 55 Hz for the transfer certificate.

We report macro-F1 for SEED and Sleep-EDF and macro-averaged AUROC for PTB-XL. To quantify sampling-rate sensitivity, we calculate the score drop relative to each model’s training-rate score. The worst drop is the largest decrease across its target rates, and we average this quantity over the trained models of each design. Reporting both the training-rate score and the mean worst drop allows us to assess classification performance together with transfer stability.

### 5.2. Baselines and implementation details

All designs use the backbone described in Section 3.2 and differ only in the first temporal convolutional layer. This layer contains sixteen filters with a duration of 250 ms. ProlateNet uses a bandwidth of 55 Hz and twenty-five basis functions, following the design rules in Section 4. All continuous kernels are evaluated at exact physical time lags with the same endpoint weighting, so their comparison isolates differences in kernel representation.

We compare fourteen first-layer designs, including ProlateNet. The discrete CNN baseline retains its learned taps at every target rate. General continuous kernels include Sinc [17], natural cubic splines with different numbers of control points, and a sinusoidal representation network (SIREN) corresponding to the kernel generator used in CKConv [12]. We also implement FlexConv [13] and sampling-frequency-independent layers [11] using the published descriptions and authors’ code.

To examine the role of spectral concentration, we include controls with constrained frequency ranges. FlexConv-fit initializes the kernel within its training-Nyquist anti-aliasing constraint, while SFI-band places the filter center frequencies within the signal band. Additional variants use the same 55-Hz boundary as ProlateNet. A cosine-basis control also matches ProlateNet in both nominal band-width and number of basis functions, allowing us to examine how the basis choice affects transfer.

We additionally evaluate multi-rate augmentation and two alternative deployment strategies. For augmentation, the CNN and Spline-31 are trained on mini-batches resampled to different rates. For input conversion, CNN + FIR resampling converts target-rate recordings to the CNN’s training rate before inference. For adaptation after training, prolate completion projects the learned CNN taps onto a continuous prolate kernel space.

We select learning rates using the validation partitions and report the remaining architecture and optimization settings in Supplementary Table S1. Within each dataset, the designs use the same evaluation partitions and random-seed sets. These shared settings allow paired comparisons of training-rate performance and sampling-rate sensitivity.

We define stability as a mean worst drop below 0.01. A design is accurate when its training-rate score is not significantly lower than the CNN’s in a paired two-sided test at a significance level of 0.05, and it meets the joint comparison criterion when it is both stable and accurate. For the EEG datasets, we use a Wilcoxon signed-rank test on fold-level scores averaged over random seeds. PTB-XL has a single official test fold, so we use a paired bootstrap over its 2,158 test records, with each design’s score averaged over its five seeds. The complete paired comparisons are reported in the Supplementary Material.

### 5.3. Comparison with continuous and discrete kernels

ProlateNet maintains stable transfer across all three datasets. Among the fourteen single-rate first-layer designs, ProlateNet and Spline-15 meet the joint comparison criterion in all four settings of Table 2. ProlateNet’s mean worst drop remains below 0.003 throughout these comparisons. By contrast, the discrete CNN exhibits mean worst drops ranging from 0.21 to 0.40, as shown in Fig. 2.

**Table 2.** Score at the training rate / mean worst drop over all target rates. Every continuous kernel is sampled at the exact lags *n*/*f* with trapezoid end weights. Higher scores and smaller drops are better. Bold cells meet both criteria of Section 5.2, a mean worst drop below 0.01 and a training-rate score not significantly below that of the CNN, and are marked for the single-rate first layers only. Multirate augmentation changes the training data, and the reference rows change the input or the trained taps. Supplementary Section S11 lists the paired tests behind every bold cell.

| Design | SEED, macro-F1 | Sleep-EDF, macro-F1 | PTB-XL, 100 Hz, macro-AU-ROC | PTB-XL, 500 Hz, macro-AU-ROC |
|---|---|---|---|---|
| ProlateNet | **0.527 / 0.0026** | **0.714 / 0.0020** | **0.892 / 0.0003** | **0.892 / 0.0001** |
| CNN | 0.546 / 0.3225 | 0.714 / 0.4026 | 0.891 / 0.3042 | 0.889 / 0.2069 |
| Spline-15 | **0.544 / 0.0053** | **0.713 / 0.0036** | **0.893 / 0.0004** | **0.891 / 0.0007** |
| Spline-31 | 0.528 / 0.0120 | **0.710 / 0.0039** | **0.891 / 0.0004** | **0.893 / 0.0016** |
| Spline-63 | 0.542 / 0.1341 | 0.712 / 0.5399 | 0.892 / 0.1130 | 0.893 / 0.0780 |
| Sinc | **0.525 / 0.0006** | 0.689 / 0.0002 | 0.882 / −0.0001 | 0.883 / 0.0001 |
| SIREN | 0.494 / 0.1356 | 0.582 / 0.1154 | 0.856 / 0.0198 | 0.856 / 0.0191 |
| Cosine basis | 0.532 / 0.0151 | 0.715 / 0.0169 | **0.892 / 0.0039** | **0.892 / 0.0027** |
| FlexConv | 0.527 / 0.1780 | 0.699 / 0.0231 | 0.888 / 0.0052 | 0.888 / 0.0334 |
| FlexConv-fit | 0.516 / 0.0054 | 0.693 / 0.0256 | 0.882 / 0.0025 | 0.881 / 0.0365 |
| FlexConv-fit (55 Hz) | 0.516 / 0.0136 | 0.700 / 0.0473 | 0.883 / 0.0021 | 0.879 / 0.0040 |
| SFI | 0.536 / 0.3522 | 0.709 / 0.0894 | 0.882 / 0.0603 | 0.871 / 0.2636 |
| SFI-band | 0.522 / 0.0039 | 0.710 / 0.0450 | 0.882 / 0.0331 | 0.880 / 0.0032 |
| SFI-band (55 Hz) | 0.532 / 0.2233 | 0.705 / 0.1229 | 0.883 / 0.0291 | 0.879 / 0.1417 |
| *Multi-rate augmentation* |  |  |  |  |
| CNN | 0.523 / 0.0276 | 0.683 / 0.0368 | 0.870 / 0.0148 | 0.859 / −0.0050 |
| Spline-31 | 0.525 / 0.0077 | 0.712 / 0.0028 | 0.892 / 0.0004 | 0.892 / 0.0003 |
| *Reference* |  |  |  |  |
| CNN + FIR resampling | 0.546 / 0.0019 | 0.714 / 0.0001 | 0.891 / 0.0002 | 0.889 / 0.0000 |
| CNN + prolate completion | 0.547 / 0.0089 | 0.714 / 0.0016 | 0.891 / 0.0012 | 0.889 / 0.0009 |

**Fig. 2.**
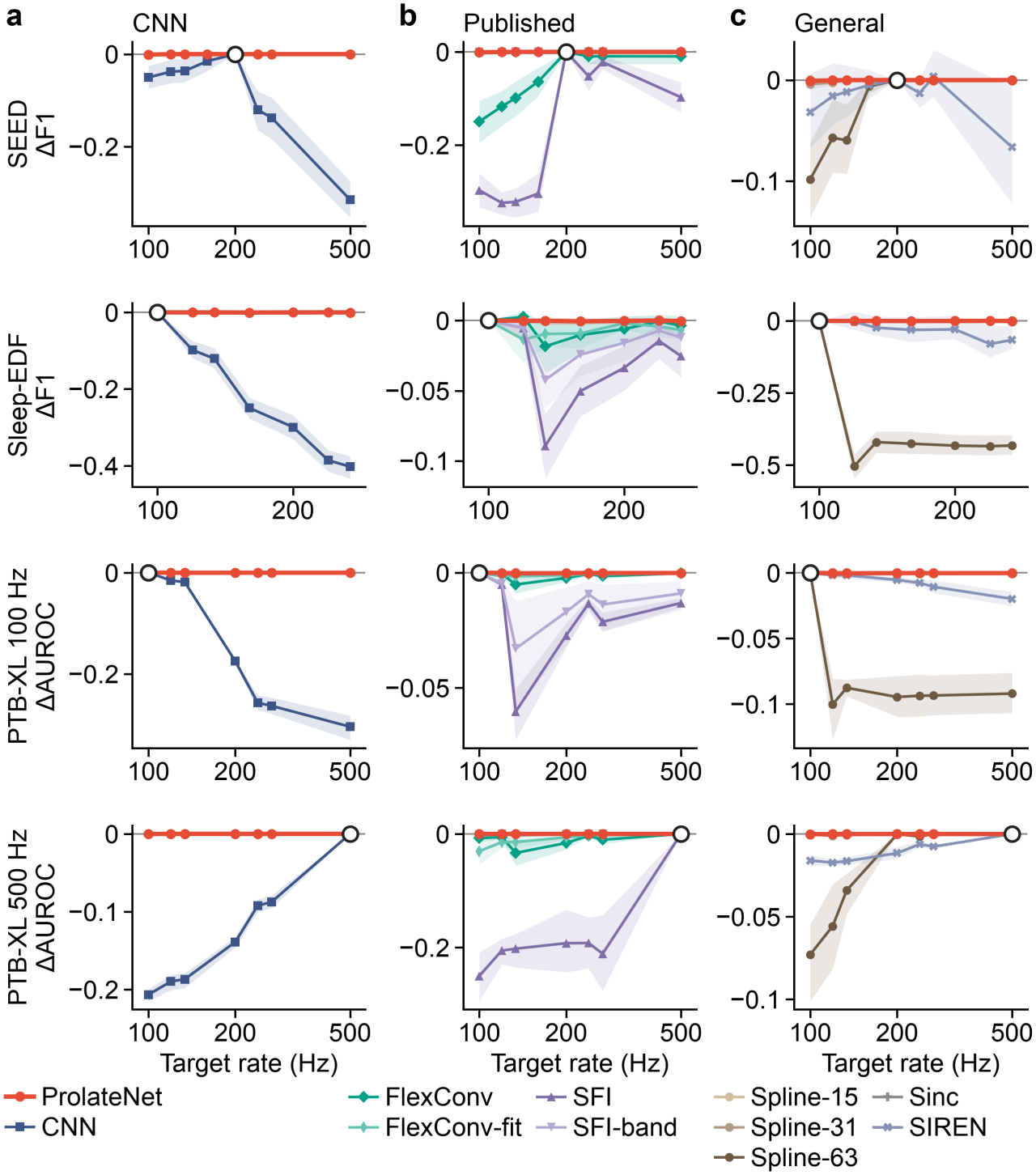
ProlateNet keeps its training-rate score at every target rate on all three datasets. Each panel shows the score change *S*(*f*) − S(*f*_0_) at target rate *f*, where *f*_0_ is the training rate (open circle). Scores are macro-F1 on SEED and Sleep-EDF and AUROC on PTB-XL, which was trained once at 100 Hz and once at 500 Hz. (a) ProlateNet and the discrete CNN. (b) ProlateNet and the published continuous-kernel layers. (c) ProlateNet and the general continuous kernels. Lines are means over models, and shaded bands are 95% bootstrap intervals over folds (SEED, Sleep-EDF) or seeds (PTB-XL). Every continuous kernel is sampled at the exact lags *n*/*f* with trapezoid end weights.

This transfer stability is accompanied by competitive performance at the training rate. On SEED, ProlateNet trails the CNN by 0.019 macro-F1. The two models have closely matched training-rate scores on Sleep-EDF. On PTB-XL, ProlateNet scores slightly above the CNN at both training rates.

Spline-15 also transfers stably under the shared exact-lag realization. ProlateNet additionally controls the entire kernel space before training: its worst-case out-of-band energy fraction stays below 1%, compared with 22% for the Spline-15 space. The cosine control further demonstrates the importance of basis selection. Despite matching ProlateNet’s nominal bandwidth and number of basis functions, it exceeds the stability threshold on both EEG datasets. On SEED, its trained kernels place approximately 9% of their energy beyond the common frequency boundary, compared with 0.05% for ProlateNet.

The published continuous-kernel layers show varying transfer performance across datasets. With their published initializations, FlexConv and SFI exhibit mean worst drops of 0.178 and 0.352 on SEED, respectively. Their frequency-constrained variants improve transfer on SEED but do not meet the joint criterion across all settings. Among the general continuous kernels, Sinc remains stable but has a lower training-rate score on Sleep-EDF and PTB-XL, while SIREN exceeds the stability threshold on both EEG datasets. These comparisons show the importance of evaluating transfer stability together with task performance.

Multi-rate augmentation improves some designs, but its effect depends on the kernel representation. The augmented CNN retains mean worst drops more than ten times those of ProlateNet on both EEG datasets. Augmentation makes Spline-31 stable on SEED, accompanied by a significant reduction in its training-rate score relative to the CNN. The reference strategies also provide effective transfer: input resampling keeps the CNN stable, and post-training prolate completion brings its mean worst drop below the stability threshold on every dataset.

Fig. 3 illustrates the trained kernel spectra, which provide a link between these performance differences and the error analysis. Larger spline spaces and SIREN can retain substantial energy beyond the common frequency boundary, while ProlateNet concentrates its energy within the prescribed band. This motivates the next analysis: whether the spectral quantity calculated from trained weights can rank designs by their measured transfer performance.

**Fig. 3.**
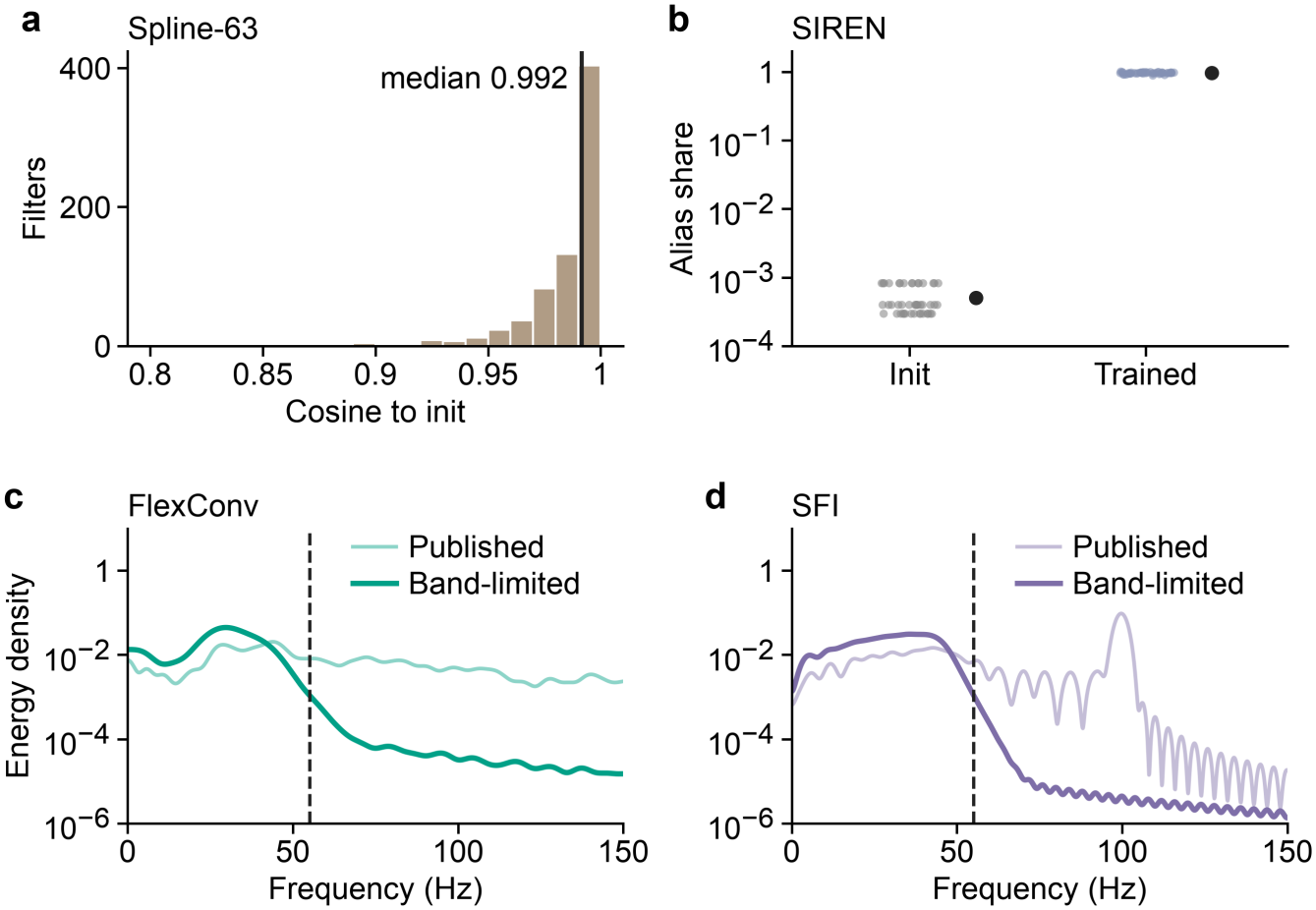
The failing continuous kernels place their energy above *f*_min_ − *B*. (a) Spline-63: cosine between the kernel component that the 200-Hz grid barely sees and the same component at initialization, over all filters; the line marks the median. (b) SIREN: share of the in-band response at 200 Hz that exists only through aliasing, at initialization and after training; dots are models, black markers the mean with a 95% bootstrap interval. Panels a and b use the Spline-63 and SIREN models of Table 2. (c, d) Energy spectra of trained FlexConv (c) and SFI (d) kernels with the published initialization and with the bandlimited initialization, averaged over models; the dashed line marks *f*_min_ − *B* = 55 Hz.

### 5.4 The transfer certificate predicts transfer from the weights

We next asked whether these failures can be read from the trained weights before any data at the target rate exist, and the transfer certificate does this. Over the 38 continuous designs trained at a single rate on SEED, its rank correlation with the measured worst drop is ρ = 0.918. Energy beyond the training Nyquist frequency *f*_0_/2, the quantity that FlexConv bounds, ranks the same designs with ρ = 0.675 only, so the ranking depends on placing the threshold at *f*_min_ − *B*. Fig. 4 plots the ranking, and Table 3 gives its bootstrap interval and the correlation for each group of designs.

**Table 3.** Rank correlation between the transfer certificate and the measured worst drop, with 95% bootstrap intervals over designs.

| Dataset | Designs or models | n | Spearman ρ [95% CI] |
|---|---|---|---|
| SEED | all single-rate continuous de-signs | 38 | 0.918 [0.794, 0.974] |
| SEED | without the published layers and their 55-Hz variants | 32 | 0.966 [0.907, 0.984] |
| SEED | prolate designs with *W* ≤ 55 Hz | 14 | 0.965 [0.838, 0.995] |
| SEED | prolate designs with *W* > 55 Hz | 12 | 0.958 [0.772, 1.000] |
| SEED | single models | 1,710 | 0.644 |
| Sleep-EDF | all single-rate continuous de-signs | 79 | 0.876 [0.813, 0.915] |
| Sleep-EDF | without the published layers and their 55-Hz variants | 73 | 0.901 [0.828, 0.939] |
| Sleep-EDF | prolate designs with *W* ≤ 55 Hz | 34 | 0.956 [0.883, 0.985] |
| Sleep-EDF | prolate designs with *W* > 55 Hz | 33 | 0.911 [0.769, 0.966] |
| Sleep-EDF | single models | 2,370 | 0.546 |

**Fig. 4.**
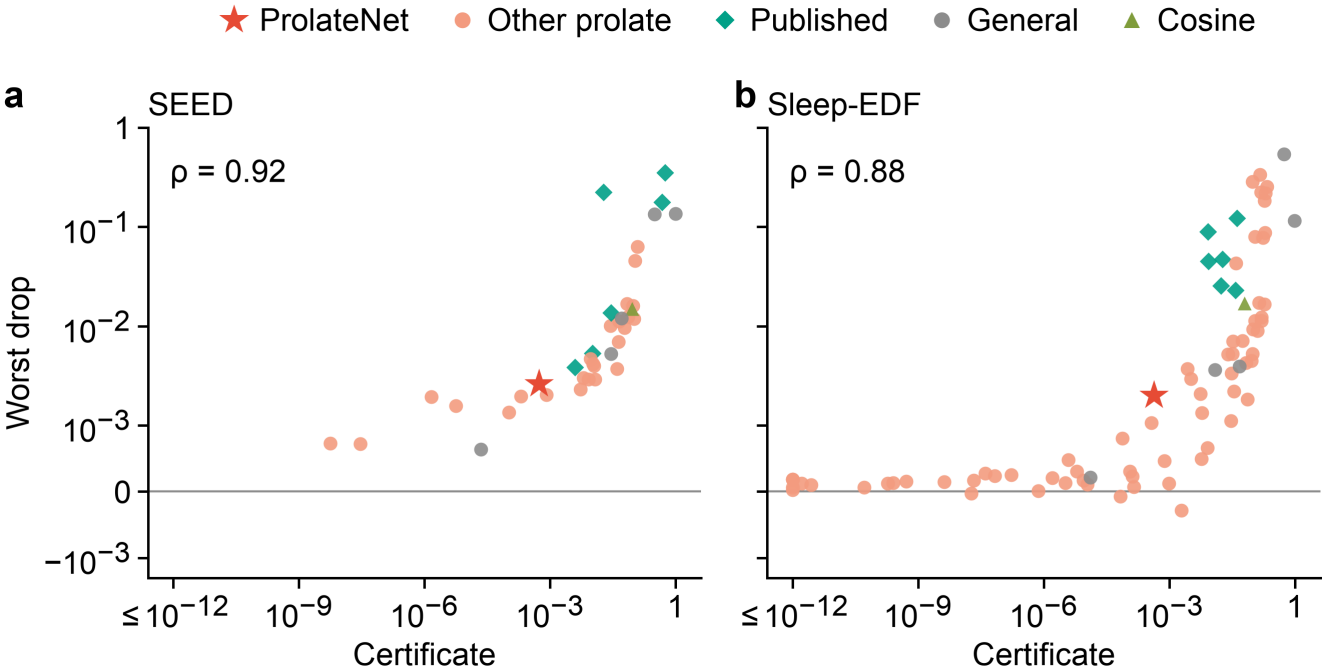
The transfer certificate ranks designs by their worst drop. Each point is a design trained at a single rate: its certificate, the kernel energy share above *f*_min_ − *B* averaged over models, against its mean worst drop over the target rates; certificates below 10^− 12^ are drawn at 10^− 12^. (a) SEED, 38 continuous designs, the four published layers and their 55-Hz variants among them, ρ = 0.918. (b) Sleep-EDF, 79 designs, ρ = 0.876. Over single models the correlation is 0.644 on SEED and 0.546 on Sleep-EDF.

The certificate also separates the published layers without a single test at the target rate. With their published initialization, FlexConv and SFI carry 47% and 55% of their kernel energy above *f*_min_ − *B*. Fig. 3c and d show where this energy lies, and these are the two published designs that fail on SEED. Their band-limited variants carry at most 1%, and both transfer. A practitioner who receives such a trained layer can therefore bound its first-layer alias error against *f*_min_ − *B* before deployment.

On Sleep-EDF the certificate ranks 79 designs with ρ = 0.876, and the prolate designs whose band stays within *f*_min_ − *B* even more closely, as Table 3 shows. The published layers are the exception on this dataset, where the four published variants carry certificates of at most 4% and the certificate does not order their drops. SFI carries the smallest certificate of the four, yet it loses the most, 0.089 F1. The certificate bounds only the aliased part of the first-layer error.

### 5.5. Parameter sensitivity: number of basis functions and band

Theory predicts where rate transfer breaks down as the kernel grows, and our data confirm that the rule *W* = *f*_min_ − *B* gives the best band. On Sleep-EDF we trained 330 models in each of six bands from 45 to 75 Hz, with *P* from 12 to 32. On SEED we added four bands around the main one, with 180 models each. As *P* grows, transfer breaks down in two stages, an onset and a collapse that Table 4 defines and Fig. 5 plots.

**Table 4.** Onset and collapse of rate transfer. Onset: *P* at which the largest absolute change of macro-F1 over the target rates first reaches 0.01, with 95% bootstrap interval over folds and seeds; the difference to the onset at 55 Hz uses the same, paired resamples. Collapse: *P* at which the mean worst drop first reaches 0.1. “>” marks a crossing beyond the largest *P* trained, 32 on Sleep-EDF and 28 on SEED (32 at 55 Hz), and “–” an interval end beyond it.

| Dataset | *W* (Hz) | 2DW | Onset *P*<br>[95% CI] | Onset − onset<br>at 55 Hz<br>[95% CI] | Collapse<br>*P* |
|---|---|---|---|---|---|
| Sleep-EDF | 45 | 22.5 | 24.0<br>[23.0, 24.2] | −4.0<br>[−5.0, −3.4] | 26.5 |
| Sleep-EDF | 50 | 25.0 | 26.1<br>[25.4, 26.1] | −2.0<br>[−2.6, −1.3] | 28.3 |
| Sleep-EDF | 55 =<br>*f*<sub>min</sub> − *B* | 27.5 | 28.0<br>[27.3, 28.2] | — | 30.4 |
| Sleep-EDF | 60 | 30.0 | 27.3<br>[26.2, 29.2] | −0.7<br>[−1.9, +1.3] | >32 |
| Sleep-EDF | 65 | 32.5 | 26.1<br>[24.2, 28.8] | −1.9<br>[−3.8, +0.8] | >32 |
| Sleep-EDF | 75 | 37.5 | 22.2<br>[20.8, 26.9] | −5.8<br>[−7.2, −1.1] | >32 |
| SEED | 50 | 25.0 | 25.8<br>[25.3, 26.3] | −1.8<br>[−2.7, −0.8] | >28 |
| SEED | 55 =<br>*f*<sub>min</sub> − *B* | 27.5 | 27.6<br>[26.6, 28.3] | — | >32 |
| SEED | 60 | 30.0 | 26.9<br>[25.5, –] | −0.7<br>[−1.9, –] | >28 |
| SEED | 65 | 32.5 | 23.9<br>[22.5, 27.9] | −3.7<br>[−5.0, +0.1] | >28 |
| SEED | 75 | 37.5 | 19.5<br>[18.5, 22.5] | −8.1<br>[−8.8, −5.6] | >28 |

**Fig. 5.**
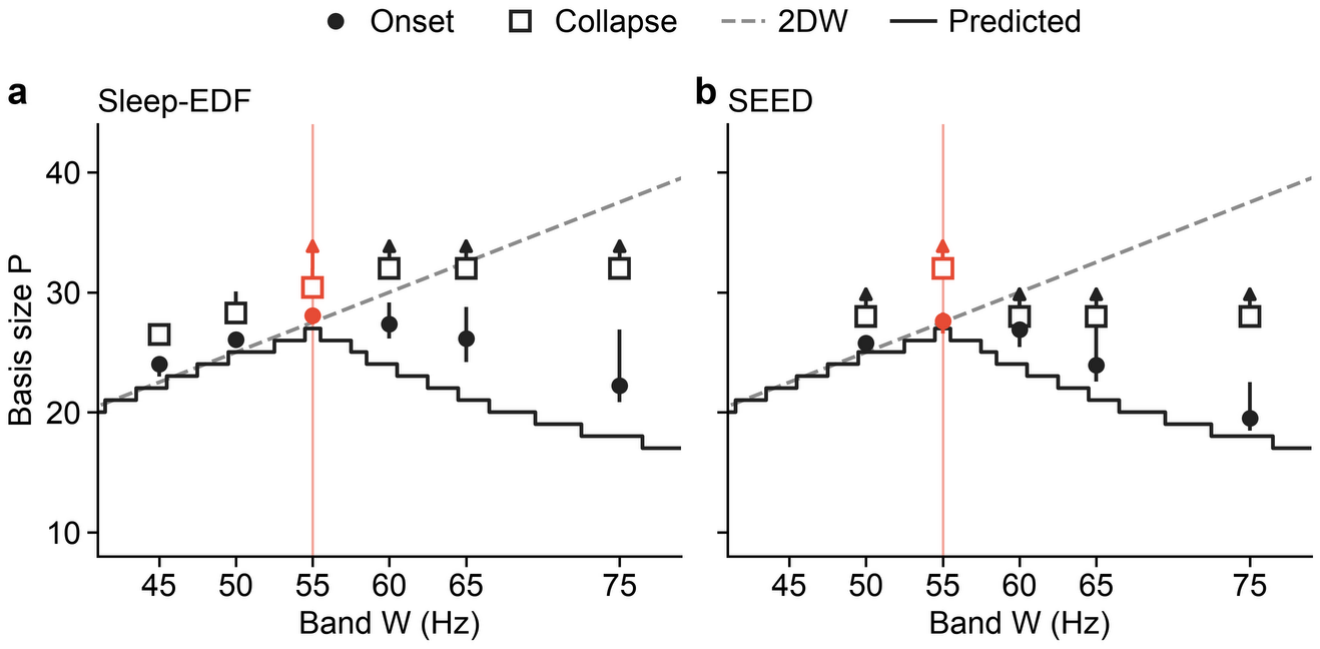
Rate transfer degrades latest at *W* = *f*_min_ − *B*. Onset (filled; *P* at which the largest |ΔF1| over the target rates reaches 0.01) and collapse (open; *P* at which the mean worst drop reaches 0.1) against the band *W*, with 95% bootstrap intervals over folds and seeds, for (a) Sleep-EDF, 330 models per band, and (b) SEED, 180 models per band. An arrow marks an estimate or interval end beyond the largest *P* trained. The dashed line is 2DW; the solid curve is the smallest *P* whose worst-case certificate exceeds 0.05 (Proposition 2(ii)), which peaks at *W* = 55 Hz with *P* = 27.

The onset comes latest at *W* = *f*_min_ − *B* on both datasets, at *P* = 28.0 on Sleep-EDF and 27.6 on SEED. Every narrower and every wider band starts earlier. The gap is significant at 45, 50 and 75 Hz on Sleep-EDF and at 50 and 75 Hz on SEED, whereas at 60 and 65 Hz its interval includes zero. Proposition 2 predicts this peak, because no band keeps more basis functions below a given certificate than *f*_min_ − *B*, and our main setting *P* = 25 sits below the onset on both datasets. On Sleep-EDF, collapse follows 2DW. For *W* ≤ 55 Hz it sets in three to four basis functions above 2DW, and for the wider bands, as for every band on SEED, it lies beyond the largest *P* we trained.

The rule also removes the need to tune *W*. When we choose *W* inside each SEED training fold, the selected models do not score above the rule. A narrower 40-Hz band scores higher at the training rate, though not significantly, and its worst drop rises to 0.0042.

### 5.6. Ablation of the realization rule

Exact-lag sampling is the component that keeps the deployed taps faithful to the trained kernel. To isolate it, we kept every trained coefficient, gain and head of the 45 SEED models and replaced only the tap rule. Table 5 lists the three alternatives, among them the linspace taps of the spline, Sinc and SIREN layers.

**Table 5.** Realization-rule ablation on SEED, 45 models per setting. Only the tap rule of the trained models changes. Response change: worst in-band change of the realized response between 200 Hz and a target rate, mean over models.

|  | *W* = 55 Hz, *P* = 25 |  |  | *W* = 45 Hz, *P* = 22 |  |  |
|---|---|---|---|---|---|---|
| Tap rule | Worst drop | p | Response change | Worst drop | p | Response change |
| Exact lags, trapezoid end weights (deployed) | 0.0026 | — | 0.068 | 0.0023 | — | 0.052 |
| Linspace taps | 0.0207 | 0.018 | 0.667 | 0.0148 | 0.0006 | 0.552 |
| Linspace taps, trape-zoid end weights | 0.0213 | 0.0054 | 0.673 | 0.0163 | 0.0076 | 0.551 |
| Exact lags, no end weights | 0.0022 | 0.026 | 0.073 | 0.0049 | 0.012 | 0.040 |

p: fold-level Wilcoxon signed-rank test of the worst drop against the deployed rule, 15 folds.

Linspace taps raise the worst drop of the main ProlateNet setting about eight times, as Table 5 and Fig. 6a show. The weights reveal the same change before any data are used. Linspace taps depart from the spacing 1/*f* and compress the kernel in time at 100, 125, 250 and 500 Hz. The worst in-band response change between 200 Hz and a target rate grows accordingly, about ten times. Trapezoid end weights matter little, since removing them moves the worst drop by less than 0.003, in opposite directions for the two settings.

**Fig. 6.**
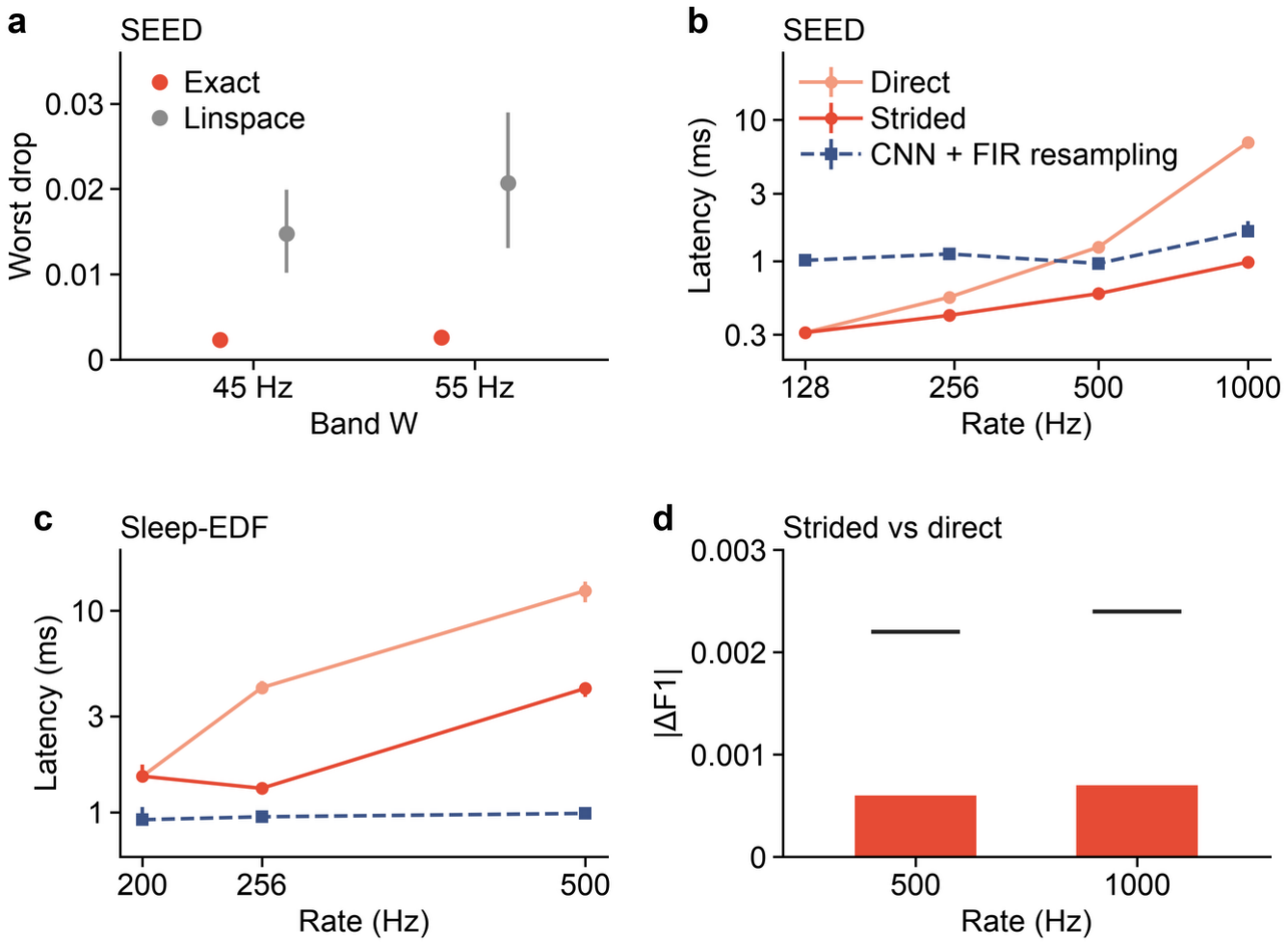
Exact lags and strided output keep deployment stable and cheap. (a) SEED worst drop with exact lags and with linspace taps for the 55-Hz and 45-Hz bases; means over 45 models with 95% bootstrap intervals. (b, c) Latency per window on one CPU core for the direct and strided ProlateNet paths and for CNN + FIR resampling, median with interquartile range over 1,000 calls; the strided path equals the direct path at the lowest rate. (d) Mean |ΔF1| between the strided and the direct path on SEED at 500 Hz (*s* = 4) and 1000 Hz (*s* = 8); ticks mark the largest value over 45 models.

### 5.7. Complexity and runtime

ProlateNet runs directly at the target rate, so its first layer keeps 400 parameters at every rate, within 6,547 for the whole SEED network. It also avoids the 50 to 100 ms group delay that a linear-phase polyphase resampler adds at these rates. A new target rate costs only a one-time tap build of under 1 ms and about 40 KiB of taps and realization matrix.

Above the training rate, the strided output of Section 4.6 keeps the first layer cheap. We tested it on SEED at 1000 Hz, where *s* = 8. There it cuts the arithmetic of the first layer per window about eight times, and Fig. 6d shows that window macro-F1 changes by at most 0.0024. The error does not grow with the stride, since the 500-Hz grid with half that stride gives changes of the same size.

To compare deployment cost, we measured latencies on one CPU core at batch size 1, as Table 6 and Fig. 6b and c report. Below the training rate, ProlateNet is more than three times faster than CNN + FIR resampling. Above it, the strided path also beats CNN + FIR at every rate on SEED, whereas on the two-channel Sleep-EDF recordings CNN + FIR is faster than both ProlateNet paths. Every median latency stays below 13 ms, far shorter than the analysis windows of Table 1.

**Table 6.** Deployment cost per window on one CPU core (Apple M4 Pro, float32, batch 1, one thread), median latency in ms over 1,000 calls from one session. s: output stride. FIR delay: group delay of the linear-phase FIR used by CNN + FIR, which ProlateNet does not need.

| Data | *f* (Hz) | ProlateNet direct | ProlateNet strided (s) | CNN + FIR resampling | FIR delay (ms) |
|---|---|---|---|---|---|
| SEED | 128 | 0.31 | = direct (1) | 1.01 | 78 |
| SEED | 250 | 0.55 | 0.41 (2) | 1.12 | 50 |
| SEED | 500 | 1.25 | 0.59 (4) | 0.96 | 50 |
| SEED | 1000 | 6.90 | 0.98 (8) | 1.62 | 50 |
| Sleep-EDF | 200 | 1.51 | = direct (1) | 0.92 | 100 |
| Sleep-EDF | 256 | 4.15 | 1.31 (2) | 0.95 | 100 |
| Sleep-EDF | 500 | 12.55 | 4.12 (4) | 0.99 | 100 |

## 6. Conclusion

We have presented ProlateNet, a framework for transferring convolutional networks across sampling rates without retraining. We derive a uniform bound on offset-averaged first-layer aliasing error and use it to guide kernel bandwidth and basis size. The resulting design combines prolate spheroidal wave function kernels, exact-lag sampling and a transfer certificate calculated from trained weights. Experiments on three public EEG and ECG datasets demonstrate stable transfer with competitive performance at the training rate. Across these datasets, the mean worst drop at unseen rates remains below 0.003. Comparisons across fourteen first-layer designs and matched controls show the importance of spectral concentration and faithful kernel realization. The certificate also ranks designs by measured transfer performance, connecting the theoretical analysis with empirical behavior.

ProlateNet provides a practical approach to reusing physiological signal models when recording rates change. Its compact parameterization keeps the number of learned kernel coefficients fixed as the deployment sampling rate increases. We use the design rules to determine the kernel space before training and the certificate to assess trained kernels before deployment. We thereby connect model construction, learning and deployment through a common spectral criterion. Researchers can also use this criterion to investigate the transfer properties of other continuous-kernel representations. Here, we consider uniformly sampled, bandlimited inputs and concentrate rate-dependent processing in the first temporal layer. In future work, we will extend the analysis to nonuniform sampling and deeper networks with multiple rate-dependent layers and nonlinear activations. We will use these extensions to address a wider range of acquisition settings and network architectures.

## Ethics statement

This study is based solely on the previously collected SEED, Sleep-EDF and PTB-XL datasets, obtained under the dataset providers’ terms, and no new data from human participants were collected.

## CRediT authorship contribution statement

Han Cui: Conceptualization, Methodology, Software, Validation, Formal analysis, Investigation, Data curation, Visualization, Writing – original draft. Yuge Zhu: Validation, Writing – review & editing. Bing Li: Supervision, Writing – review & editing. Shaojia Huang: Supervision, Funding acquisition, Writing – review & editing. Tao Xu: Supervision, Project administration, Resources, Funding acquisition, Writing – review & editing.

## Data availability

SEED is available on request from the BCMI laboratory of Shanghai Jiao Tong University under its data-use licence, which does not permit redistribution. Sleep-EDF and PTB-XL are openly available on PhysioNet. The results generated in this study will be made publicly available upon journal publication.

## Code availability

The code for all analyses and figures will be made publicly available upon journal publication.

## Declaration of generative AI and AI-assisted technologies in the manuscript preparation process

During the preparation of this work the authors used Claude (Anthropic) in order to assist with code development and to improve the readability of the manuscript.

After using this tool, the authors reviewed and edited the content as needed and take full responsibility for the content of the published article.

## Appendix A. Proofs of Propositions 1–3

This appendix proves the three propositions of Section 4. The Supplementary Material gives the rate-transfer identity (3) with its proof, the spectrum-weighted and two-rate forms of Proposition 1, the fixed-grid analysis, the realization rank, the cascade bound and the numerical checks.

### A.1 Setting and notation

We write *T* = *D*/2 and *I* = [−*T, T*]. A kernel is a function *k* ∈ *L*^2^(ℝ) that vanishes outside *I*; the fixed gain *g*_*j*_ of (7) is absorbed into *k*. Its spectrum *K*(*ξ*) = *∫*ℝ *k* (*t*)*e*^−π*iξt*^ *dt* extends to an entire function because *k* has bounded support, and Plancherel’s theorem gives ∥ *K* ∥_2_=∥ *k* ∥_2_. For a measurable set Ω ⊆ ℝ we write *ℰ*_Ω_(*k*) = *∫*_Ω_ | *K*|^2^, together with *ℰ*_≥*V*_(*k*) = *ℰ*_{|_*ξ*_|≥*V*}_(*k*) and *ℰ*_X*V*_(*k*) = *ℰ*_{|S|X*V*}_(*k*). Throughout, *f*_min_ > 2*B* and *W*_0_ = *f*_min_ − *B*.

#### Offset realization

For *θ* ∈ [0,1) the *θ*-offset taps at rate *f* are *h*_*f,θ*_[*n*] = *k*((*n* + *θ*)/*f*)/*f*, placed at the physical lags *t*_*n*_ = (*n* + *θ*)/*f*, and the response referred to physical time is 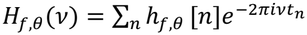. The case *θ* = 0 is (1)– (2). For fixed *n*, changing *k* on a null set changes *h*_*f,θ*_[*n*] only for a null set of *θ*, and an end lag *t*_*n*_ = ±*T*, where the weight *w*_*n*_ of (11) acts, occurs for at most two values of *θ*. The error (4) therefore depends on *k* only as an element of *L*^2^(*I*).

**Alias band**. Let *A*_*f*_ = ⋃_*m≠* 0_(*mf* + [−*B, B*]). For |*v*| ≤ *B* and *m* ≠ 0 we have |*mf* + *v*| ≥ |*m*|*f* − *B* ≥ *f* − *B*, hence

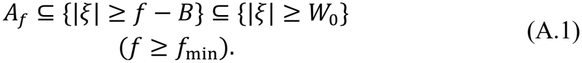

**Prolate functions**. Let *Q* be the operator (8) on *L*^2^(*I*) with *W* = *W*_0_. Plancherel’s theorem gives 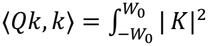, so that

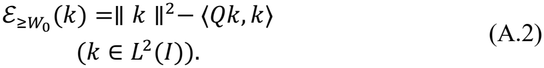

The operator *Q* is compact, self-adjoint and positive. It is also injective: ⟨*Qk, k*⟩ = 0 forces *K* = 0 on (−*W*_0_, *W*_0_), hence *K* ≡ 0 because *K* is entire, and *k* = 0. By the spectral theorem, its eigenfunctions *φ*_*p*_ = *φ*_*p*_(⋅; *D, W*_0_) form an orthonormal basis of *L*^2^(*I*), with simple eigenvalues 1 > *λ*_5_ > *λ*_)_ > … > 0 that tend to 0 [16]. For *k* = ∑_*p*_ *a*_*p*_ *φ*_*p*_, (A.2) gives

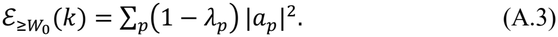

We write *λ*_*p*_ = *λ*_*p*_(*D, W*_0_), *N*_*α*_ (*D, W*_0_) = #{*p*: *λ*_*p*_ ≥ *α*}, which is finite for *α* > 0, and *V*_*p*_(*D, W*) = span{*φ*_1_(⋅; *D, W*), …, *φ*_*p*_(⋅; *D, W*)} for the PKC space with band *W* and size *P*. For a subspace *V*⊆ *L*^2^(*I*), the worst-case certificate is 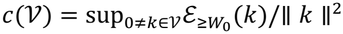.

### A.2. Proof of Proposition 1

Proposition 1 holds for every kernel *k* ∈ *L*^2^(*I*); bounded variation is needed only for the pointwise identity (3) (Supplementary Theorem S3.1).

*Step 1 (Fourier coefficients in the offset)*. Fix *f* and *v*. Only finitely many *n* give lags *t*_*n*_ ∈ *I* for some *θ* ∈ [0,1). For each of them the substitution *t* = (*n* + *θ*)/*f* gives 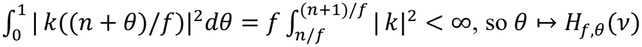 belongs to *L*^2^(0,1). With the same substitution, and *e* ^− 2*πimθ*^ = *e*^− 2*πimft*^ because *e*^− 2*πimn*^= 1,

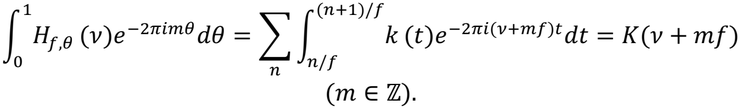

*Step 2 (Parseval’s identity)*. The function *θ ↦ H*_*f,θ*_(*v*) − *K*(*v*) has the Fourier coefficients *K*(*v* + *mf*) for *m* ≠ 0 and 0 for *m* = 0. Parseval’s identity in *L*^2^(0,1) gives

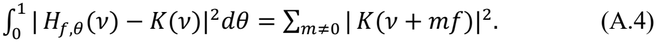

*Step 3 (integration over the signal band)*. The integrand of (4) is nonnegative and jointly measurable in (*θ, v*). Tonelli’s theorem and (A.4) give

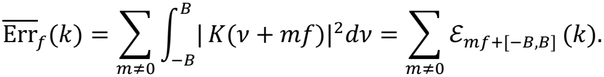

For *f* ≥ *f*_min_ > 2*B*, consecutive intervals *mf* + [−*B, B*] are separated by gaps of length *f* − 2*B* > 0, so the intervals are pairwise disjoint and the sum equals 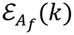. This is the equality in (5). Since *ℰ*_Ω_ is monotone in Ω, the two inequalities in (5) follow from (A.1).

*Step 4 (near-attainment)*. For *f* = *f*_min_ the bands *m* = ±1 of 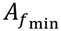 are ±[*W*_0_, *f*_min_ + *B*]. Hence 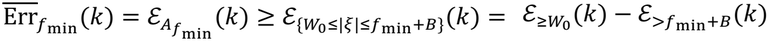. □

Averaging over *θ* makes the alias orders orthogonal through the characters *e*^2*πimθ*^, which is why their energies add in (A.4); with *S* ≡ 1 and restricted to the signal band, the resulting equality is the shift-averaged error formula of Blu and Unser [37].

### A.3 Proof of Proposition 2

We prove the two statements of Proposition 2 in the following form.

i. Every *P*-dimensional subspace *V*⊆ *L*^2^(*I*) satisfies *c*(*V*) ≥ 1 – *λ*_*p*_, with equality for *V*= *V*_B_(*D, W*_0_).
ii. For 0 < *ε* < 1 and *W* > 0, let *P*_*ε*_(*W*) be the largest *P* ≥ 1 with *c*(*V*_*P*_(*D, W*)) ≤ *ε*, and *P*_*ε*_(*W*) = 0 if no such *P* exists. Then *P ε* (*W*) ≤ *N*_1−_ *ε* (*D, W*_0_) = *P ε* (*W*_0_) for every *W* > 0.

*Proof*. (i) The linear map *k* ↦ (⟨*k, φ*_1_⟩, …, ⟨*k, φ*_*P*−1_⟩) from *V*to ℂ ^*P*−1^ has a non-trivial null space, so *V*contains some *k* ≠ 0 with *a*_1_ = … = *a*_*p*−1_ = 0. For this kernel ⟨*Qk, k*⟩ = ∑_*p*≥*P*_ *λ*_*p*_ |*a*_*p*_|^)^ ≤ *λ*_*P*_ ∥ *k* ∥^2^, and (A.2) gives 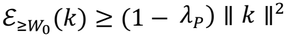. Hence *c*(*V*) ≥ 1 – *λ*_*P*_. For *V*= *V*_*P*_(*D, W*_0_), every *k* = ∑_*p*≤ *P*_ *a*_*p*_ *φ*_*p*_ satisfies 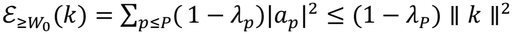 by (A.3), with equality at *k* = *φ*_*P*_. This is the Courant–Fischer principle [40,41] for *Q*.

(ii) The spaces *V*_*P*_(*D, W*) increase with *P*, and *c*(*V*_*P*_(*D, W*)) ≥ 1 − *λ*_*P*_ → 1 by (i), so *P*_*ε*_(*W*) is well defined. If *c*(*V*_*P*_(*D, W*)) ≤ *ε*, then (i) gives 1 − *λ*_*P*_ ≤ *ε*, that is, *λ*_*P*_ ≥ 1 − *ε*. Since the eigenvalues decrease strictly, {*p*: *λ*_*p*_ ≥ 1 − *ε*} = {1, …, *N*_1−*ε*_(*D, W*_0_)}, so *P* ≤ *N*_1−*ε*_(*D, W*_0_). For *W* = *W*_0_, (i) gives *c*(*V*_*P*_(*D, W*_0_)) = 1 − *λ*_*P*_, which is at most *ε* exactly when *P* ≤ *N*_1−*ε*_(*D, W*_0_). □

The certificate *c*(*V*) is a worst case over the whole space: by Proposition 1, every member *k* of *V*satisfies 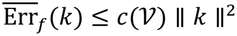 at every rate *f* ≥ *f*_min_.

### A.4 Proof of Proposition 3

For a set of rates *F* we write Ω_*F*_ = ⋃_*f*∈*F*_*A*_*f*_ for the frequencies that alias at some rate of *F*.

**Lemma A.1 (coverage)**. Let 2*B* < *f*_min_ < *f*_max_ and *F* = [*f*_min_, *f*_max_]. Then Ω_*F*_ = {|*ξ*| ≥ *W*_0_} if and only if *f*_max_ ≥ 2*f*_min_ − 2*B*. For *F* = [*f*_min_, ∞) the equality always holds.

*Proof*. The inclusion Ω_*F*_ ⊆ {|*ξ*| ≥ *W*_0_} is (A.1). For fixed *m* ≥ 1, ⋃_*f*∈*F*_(*mf* + [−*B, B*]) = [*mf*_min_ − *B, mf*_max_ + *B*], and the bands *m* ≤ −1 are the mirror images. The intervals of orders *m* and *m* + 1 meet if and only if *mf*_max_ + *B* ≥ (*m* + 1)*f*_min_ − *B*, that is, *m*(*f*_max_ − *f*_min_) ≥ *f*_min_ − 2*B*. The left side increases with *m*, so the condition holds for every *m* ≥ 1 exactly when it holds for *m* = 1, that is, when *f*_max_ ≥ 2*f*_min_ − 2*B*. The intervals then cover [*f*_min_ − *B*, ∞) = [*W*_0_, ∞). If the condition fails, the nonempty interval (*f*_max_ + *B*, 2*f*_min_ − *B*) is not covered, because the band *m* = 1 ends at *f*_max_ + *B*, the bands *m* ≥ 2 start at 2*f*_min_ − *B* or later, and the bands *m* ≤ −1 lie at negative frequencies. For *F* = [*f*_min_, ∞) the band *m* = 1 alone covers [*W*_0_, ∞). □

*Proof of Proposition 3*. Let *F* = [*f*_min_, *f*_max_] with *f*_max_ ≥ 2*f*_min_ − 2*B*, let 0 < *ε* < 1 and *N* = *N*_1−*ε*_(*D, W*_0_). By Lemma A.1, 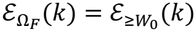 for every kernel *k*.

Let *V*⊆ *L*^2^(*I*) be a linear space with dim*V*> *N*, and choose a subspace *V*^`^ ⊆ *V*of dimension *P* = *N* + 1. Since *P* > *N* and the eigenvalues decrease, *λ*_*P*_ < 1 − *ε*, and Proposition 2(i) gives *c*(*V*^`^) ≥ 1 − *λ*_*P*_ > *ε*. By the definition of the supremum, *V*^`^ contains a kernel *k* ≠ 0 with 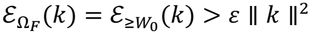. Each frequency of Ω_*F*_ lies in *A*_*f*_ for some *f* ∈ *F*, and by Proposition 1 the energy of *k* on *A*_*f*_ is its offset-averaged error at *f*.

For the attainment, let *N* ≥ 1 and *k* = ∑_*p*≤a_ *a*_*p*_ *φ*_*p*_ ∈ *V*_a_(*D, W*_0_). By (A.1) and (A.3), 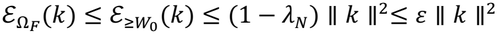, for every set of rates *F* with min*F* ≥ *f*_min_. The PKC space with *W* = *W*_0_ and *P* = *N*_1−*ε*_(*D, W*_0_) therefore attains the limit. □

The argument uses no property of the space *V*other than its dimension, so the limit applies to splines, polynomials and every other linear parameterization of kernels supported on *I*.

## References

[1] Schirrmeister RT, Springenberg JT, Fiederer LDJ, Glasstetter M, Eggensperger K, Tangermann M, Hutter F, Burgard W, Ball T. Deep learning with convolutional neural networks for EEG decoding and visualization. Hum. Brain Mapp. 38 (2017) 5391–5420.

[2] Lawhern VJ, Solon AJ, Waytowich NR, Gordon SM, Hung CP, Lance BJ. EEGNet: a compact convolutional neural network for EEG-based brain– computer interfaces. J. Neural Eng. 15 (2018) 056013.

[3] Perslev M, Darkner S, Kempfner L, Nikolic M, Jennum PJ, Igel C. U-Sleep: resilient high-frequency sleep staging. npj Digit. Med. 4 (2021) 72.

[4] Wagner P, Strodthoff N, Bousseljot R-D, Kreiseler D, Lunze FI, Samek W, Schaeffter T. PTB-XL, a large publicly available electrocardiography dataset. Sci. Data 7 (2020) 154.

[5] Kemp B, Zwinderman AH, Tuk B, Kamphuisen HAC, Oberye JJL. Analysis of a sleep-dependent neuronal feedback loop: the slow-wave microcontinuity of the EEG. IEEE Trans. Biomed. Eng. 47 (2000) 1185–1194.

[6] Goldberger AL, Amaral LAN, Glass L, Hausdorff JM, Ivanov PCh, Mark RG, Mietus JE, Moody GB, Peng C-K, Stanley HE. PhysioBank, PhysioToolkit, and PhysioNet: components of a new research resource for complex physiologic signals. Circulation 101 (2000) e215–e220.

[7] Zheng W-L, Lu B-L. Investigating critical frequency bands and channels for EEG-based emotion recognition with deep neural networks. IEEE Trans. Auton. Ment. Dev. 7 (2015) 162–175.

[8] Yang C, Westover MB, Sun J. BIOT: Biosignal transformer for cross-data learning in the wild. In: Adv. Neural Inf. Process. Syst. (NeurIPS) 36, 2023.

[9] Jiang W-B, Zhao L-M, Lu B-L. Large brain model for learning generic representations with tremendous EEG data in BCI. In: Int. Conf. Learn. Represent. (ICLR), 2024.

[10] Ellis CA, Miller RL, Calhoun VD. Cross-sampling rate transfer learning for enhanced raw EEG deep learning classifier performance in major depressive disorder diagnosis. In: Proc. IEEE Int. Symp. Biomed. Imaging (ISBI), 2024, pp. 1–5.

[11] Saito K, Nakamura T, Yatabe K, Saruwatari H. Sampling-frequency-independent convolutional layer and its application to audio source separation. IEEE/ACM Trans. Audio Speech Lang. Process. 30 (2022) 2928–2943.

[12] Romero DW, Kuzina A, Bekkers EJ, Tomczak JM, Hoogendoorn M. CKConv: Continuous kernel convolution for sequential data. In: Int. Conf. Learn. Represent. (ICLR), 2022.

[13] Romero DW, Bruintjes R-J, Tomczak JM, Bekkers EJ, Hoogendoorn M, van Gemert JC. FlexConv: Continuous kernel convolutions with differentiable kernel sizes. In: Int. Conf. Learn. Represent. (ICLR), 2022.

[14] Imamura K, Nakamura T, Takamune N, Yatabe K, Saruwatari H. Local equivariance error-based metrics for evaluating sampling-frequency-independent property of neural network. In: Proc. Eur. Signal Process. Conf. (EUSIPCO), 2025, pp. 276–280.

[15] Landau HJ, Pollak HO. Prolate spheroidal wave functions, Fourier analysis and uncertainty – III: The dimension of the space of essentially time- and band-limited signals. Bell Syst. Tech. J. 41 (1962) 1295–1336.

[16] Slepian D, Pollak HO. Prolate spheroidal wave functions, Fourier analysis and uncertainty – I. Bell Syst. Tech. J. 40 (1961) 43–63.

[17] Ravanelli M, Bengio Y. Speaker recognition from raw waveform with SincNet. In: Proc. IEEE Spoken Lang. Technol. Workshop (SLT), 2018, pp. 1021–1028.

[18] Borra D, Fantozzi S, Magosso E. Interpretable and lightweight convolutional neural network for EEG decoding: Application to movement execution and imagination. Neural Netw. 129 (2020) 55–74.

[19] Zeghidour N, Teboul O, de Chaumont Quitry F, Tagliasacchi M. LEAF: A learnable frontend for audio classification. In: Int. Conf. Learn. Represent. (ICLR), 2021.

[20] Yuan Y, Wang L-N, Zhong G, Gao W, Jiao W, Dong J, Shen B, Xia D, Xiang W. Adaptive Gabor convolutional networks. Pattern Recognit. 124 (2022) 108495.

[21] Qiu Q, Cheng X, Calderbank R, Sapiro G. DCFNet: Deep neural network with decomposed convolutional filters. In: Proc. Int. Conf. Mach. Learn. (ICML), PMLR 80, 2018, pp. 4198–4207.

[22] Saito K, Nakamura T, Yatabe K, Koizumi Y, Saruwatari H. Sampling-frequency-independent audio source separation using convolution layer based on impulse invariant method. In: Proc. Eur. Signal Process. Conf. (EU-SIPCO), 2021, pp. 321–325.

[23] Imamura K, Nakamura T, Yatabe K, Saruwatari H. Neural analog filter for sampling-frequency-independent convolutional layer. APSIPA Trans. Signal Inf. Process. 13 (2024) 1–23.

[24] Kovachki N, Li Z, Liu B, Azizzadenesheli K, Bhattacharya K, Stuart A, Anandkumar A. Neural operator: Learning maps between function spaces with applications to PDEs. J. Mach. Learn. Res. 24 (2023) 1–97.

[25] Li Z, Kovachki N, Azizzadenesheli K, Liu B, Bhattacharya K, Stuart A, Anandkumar A. Fourier neural operator for parametric partial differential equations. In: Int. Conf. Learn. Represent. (ICLR), 2021.

[26] Bartolucci F, de Bézenac E, Raonić B, Molinaro R, Mishra S, Alaifari R. Representation equivalent neural operators: a framework for alias-free operator learning. In: Adv. Neural Inf. Process. Syst. (NeurIPS) 36, 2023.

[27] Gu A, Goel K, Ré C. Efficiently modeling long sequences with structured state spaces. In: Int. Conf. Learn. Represent. (ICLR), 2022.

[28] Chen J, Song P, Zhao C. Multi-scale self-supervised representation learning with temporal alignment for multi-rate time series modeling. Pattern Recognit. 145 (2024) 109943.

[29] Sun C, Li H, Song M, Cai D, Zhang B, Hong S. Time pattern reconstruction for classification of irregularly sampled time series. Pattern Recognit. 147 (2024) 110075.

[30] Wang K, Zhu Q, Zhao J, Zheng C, Zhang D. Disentangling spatio-temporal topological representations for cross-device EEG emotion recognition. Pattern Recognit. 184 (2027) 115179.

[31] Luo G, Zheng C, Zhu L, Fan T, Tian F, Wang D, Qian K, Liu J, Sun S, Hu B. Multi-scale cross-domain and class-wise kernel discriminative alignment for EEG-based emotion recognition. Pattern Recognit. 179 (2026) 113626.

[32] Azulay A, Weiss Y. Why do deep convolutional networks generalize so poorly to small image transformations? J. Mach. Learn. Res. 20 (2019) 1–25.

[33] Zhang R. Making convolutional networks shift-invariant again. In: Proc. Int. Conf. Mach. Learn. (ICML), PMLR 97, 2019, pp. 7324–7334.

[34] Karras T, Aittala M, Laine S, Härkönen E, Hellsten J, Lehtinen J, Aila T. Alias-free generative adversarial networks. In: Adv. Neural Inf. Process. Syst. (NeurIPS) 34, 2021.

[35] Xu G, Liao W, Zhang X, Li C, He X, Wu X. Haar wavelet downsampling: A simple but effective downsampling module for semantic segmentation. Pattern Recognit. 143 (2023) 109819.

[36] Landau HJ, Widom H. Eigenvalue distribution of time and frequency limiting. J. Math. Anal. Appl. 77 (1980) 469–481.

[37] Blu T, Unser M. Quantitative Fourier analysis of approximation techniques. I. Interpolators and projectors. IEEE Trans. Signal Process. 47 (1999) 2783–2795.

[38] Jayasundara D, Zhao H, Labate D, Patel VM. PIN: Prolate spheroidal wave function-based implicit neural representations. In: Int. Conf. Learn. Represent. (ICLR), 2025.

[39] Zygmund A. Trigonometric Series, third ed. Cambridge University Press, Cambridge, 2003.

[40] Courant R. Über die Eigenwerte bei den Differentialgleichungen der mathematischen Physik. Math. Z. 7 (1920) 1–57.

[41] Fischer E. Über quadratische Formen mit reellen Koeffizienten. Monatsh. Math. Phys. 16 (1905) 234–249.

