## supplementary_information for "ProlateNet: Sampling-rate transfer of convolutional networks with a weights-only certificate for physiological signals"

Section S1 lists the training settings of every design. Sections S2–S9 state and prove the theoretical results that complement Appendix A of the main text, with a numerical check after each statement. Section S10 compares two realization rules for the general continuous kernels, and Section S11 lists the paired tests behind Table 2. Section, equation and reference numbers without the prefix S refer to the main text.

### S1. Training hyperparameters

**Supplementary Table S1.** Training settings per dataset. All designs of a dataset share these settings unless a row states otherwise. $\eta$ denotes the base learning rate; "first layer" denotes the parameters of the temporal layer (taps, spline controls, Sinc cut-offs, SIREN weights, FlexConv and SFI parameters, or prolate coefficients), and "other" denotes the spatial projection and the classifier head.

| Setting | SEED | Sleep-EDF-20 | PTB-XL |
|---|---|---|---|
| Training rate $f_0$ | 200 Hz | 100 Hz | 100 Hz and 500 Hz, trained separately |
| Input window | 4 s, 52 channels, 3 classes | 30 s, 2 derivations (Fpz-Cz, Pz-Oz), 5 stages | 10 s, 12 leads, 5 diagnostic superclasses (multi-label) |
| Selection split | per outer fold: 11 inner-training and 3 inner-validation subjects (validation subjects at cyclic offsets 1, 6 and 11 from the test subject) | per test fold: the next fold (cyclically) for validation, the remaining 8 folds for training | official folds: 1–8 training, 9 validation, 10 test |
| Optimizer | AdamW, $\beta$=(0.9,0.999), $\epsilon$=$10^{-8}$ | AdamW, $\beta$=(0.9,0.999), $\epsilon$=$10^{-8}$ | as Sleep-EDF-20 |
| Weight decay | $10^{-4}$ on all parameters | $10^{-4}$ on all parameters | $10^{-4}$ on all parameters |
| Learning-rate grid | $\eta\in\{3\times10^{-4}, 10^{-3}\}$; first layer at $10\eta$ for every continuous-kernel design, at $\eta$ for the CNN (also with multi-rate augmentation); other parameters at $\eta$ | $\eta\in\{10^{-3}, 3\times10^{-4}\}$; first layer at $10\eta$ for every design, the CNN included; other parameters at $\eta$ | as Sleep-EDF-20 |

| Setting | SEED | Sleep-EDF-20 | PTB-XL |
|---|---|---|---|
| Learning-rate selection | the rate with the higher inner-validation macro-F1 (mean over the 3 validation subjects of window macro-F1 at 200 Hz, at its best epoch); scores within $10^{-4}$ of the maximum count as ties and select $3\times10^{-4}$ | the rate with the higher validation macro-F1 at 100 Hz at its best epoch; an exact tie selects $10^{-3}$ | the rate with the higher validation macro-AU-ROC at the training rate at its best epoch; an exact tie selects $10^{-3}$ |
| Epochs and early stopping | selection fits: at most 120 epochs, stopped after 20 epochs without a validation gain above $10^{-4}$ | at most 100 epochs, stopped after 10 epochs without a validation gain above $10^{-4}$ | at most 150 epochs, stopped after 15 epochs without a validation gain above $10^{-4}$ |
| Final model | refit on all 14 training subjects from the same seed, with the selected learning rate, for the selected best epoch count | weights of the best validation epoch of the selected learning rate | as Sleep-EDF-20 |
| Batch size | 64 | 64 | 64 |
| Batches per epoch | $\lceil n/64 \rceil$, windows drawn with replacement: uniform subject, then uniform trial, then uniform window | $\lfloor n/64 \rfloor$ from a new random permutation of the training windows in every epoch | as Sleep-EDF-20 |
| Loss | cross-entropy without label smoothing | class-weighted cross-entropy, weight $n/(5\, n_c)$ for class $c$ from the training-fold counts | binary cross-entropy on the 5 superclass logits |
| Design-specific loss term | FlexConv and FlexConv-fit: anti-aliasing regularizer of FlexConv at the training grid, $\lambda$=0.1; FlexConv-fit (55 Hz): fit and regularizer at 55 Hz | as SEED | as SEED |
| Gradient clipping | global $L_2$ norm 1.0 | global $L_2$ norm 1.0 | global $L_2$ norm 1.0 |
| Input normalization | per-channel mean and population variance over the training subjects' windows at 200 Hz (11 subjects for selection fits, 14 for the refit), applied unchanged at every rate | per-channel mean and standard deviation (plus $10^{-6}$) over at most 4,096 randomly drawn training windows at the training rate, applied unchanged at every rate | as Sleep-EDF-20 |
| Head dropout | 0.2 | 0.2 | 0.2 |
| Multi-rate augmentation (CNN and Spline-31) | one rate per mini-batch, a uniform integer in 100–500 Hz; each raw window is resampled by polyphase FIR (Kaiser window, $\beta$=5) before normalization | one rate per mini-batch, drawn uniformly from the evaluation grid {100, 125, 128, 160, 200, 250, 256} Hz; each window is resampled from the training grid by polyphase FIR (Kaiser window, $\beta$=5) before normalization | as Sleep-EDF-20, over {100, 125, 128, 200, 250, 256, 500} Hz |
| Training seeds | 2024, 2025, 2026 | 2024, 2025, 2026 | 2024–2028 |

| Setting | SEED | Sleep-EDF-20 | PTB-XL |
| --- | --- | --- | --- |
| Numerics | float32, TF32 disabled, deterministic algorithms | float32, framework defaults | float32, framework defaults |

In every design the first-layer parameters run at ten times the base rate, with one exception: the SEED CNN, whose parameters all share one rate. On Sleep-EDF-20 and PTB-XL the CNN taps also run at ten times the base rate. The two pipelines also differ in how they obtain the final model. SEED refits on all training subjects for the selected epoch count, whereas Sleep-EDF-20 and PTB-XL keep the best validation checkpoint. Settings internal to the FlexConv and SFI layers follow the original papers [11,13].

### S2. Setting and notation

We mark each result below as (classical) when it restates a known theorem in our notation, and we then give its source; all other results are marked (new). When a result is classical in some parts and new in others, its status names the parts. Propositions 1–3 of the main text and Lemma A.1 are proved in Appendix A, and this supplement gives their sources and numerical checks.

**Supplementary Table S2.** Status and source of every theoretical result in this supplement.

| Result | Content | Status | Original source or proof tool |
| --- | --- | --- | --- |
| Lemma S2.1 | a realized layer filters band-limited inputs with $H_f$ | classical | Shannon (1949); Parseval's identity |
| Theorem S3.1 | rate-transfer identity $H_f(\nu)=\sum_m K(\nu+mf)$ | classical | Poisson (1823, 1827); Dirichlet (1829); Jordan (1881); Zygmund (2003) |
| Lemma S4.1 | the offset average is the mean error over all placements of the kernel on the deployment lattice | classical | change of variables; random-origin sampling of Moran (1950) |
| Theorem S4.1 | offset-averaged error = spectrum-weighted alias-band energy; lattice form; bound by the energy beyond $f_{\min}-B$ (Proposition 1 is the case $S\equiv1$ of (c), $(f)$ and (g)) | classical: (a), (b), (d), and (c) for $S\equiv1$; new: (c) with a spectrum weight, (e)–(g) | Moran (1950); L'Ecuyer & Lemieux (2000); Matheron (1970/2019); Blu & Unser (1999); Crochiere & Rabiner (1975) |
| Corollary S4.1 | main space; energy just above $f_{\min}-B$ reaches only the top of the data band | new | Theorem S4.1 |
| Proposition S5.1(a) | no fixed-grid bound with a universal constant, even under small out-of-band energy | new | explicit bumps |

| Result | Content | Status | Original source or proof tool |
|---|---|---|---|
| Proposition S5.1(b) | fixed-grid constants $\gamma_f$, $\Gamma_f$ and the computable $\Gamma_f^{\rho}$; $\gamma_f$ over 100–500 Hz | new | generalized eigenvalues |
| Proposition S5.1(c) | fixed-grid bound for kernels with $k(\pm D/2)=0$ | new | Poisson summation; Cauchy–Schwarz inequality |
| Proposition 2(i) (Appendix A.3) | PKC with $W=f_{\min}-B$ minimizes the certificate | classical | Slepian & Pollak (1961); Fischer (1905); Courant (1920) |
| Proposition 2(ii) (Appendix A.3) | the safe PKC dimension is largest at $W=f_{\min}-B$ | new (design rule) | Proposition 2(i) |
| Lemma A.1 (Appendix A.4) | coverage of the alias region by a rate interval | new | interval arithmetic |
| Proposition 3 (Appendix A.4) | capacity-invariance limit $N_{1-\varepsilon}(D,f_{\min}-B)$ | classical count, applied through the new Lemma A.1 | Proposition 2(i); Lemma A.1 |
| Counting facts (Section S7) | $N_{1/2}\in\{\lfloor 2DW\rfloor,\lfloor 2DW\rfloor+1\}$; asymptotics of $N_\alpha$ | classical | Landau (1993); Landau & Widom (1980); Landau & Pollak (1962) |
| Theorem S8.1 | full-rank realization | classical | Hogan & Lakey (2012, 2015) |
| Theorem S9.1 | cascade bound by the per-layer certificates | new | telescoping identity; Minkowski's inequality |

**Supplementary Table S3.** Symbols used in the main text and in this supplement.

| Symbol | Meaning |
|---|---|
| $t$; $\nu,\xi$ | time; frequency (Hz) |
| $f$, $f_0$, $f_{\min}$ | sampling rate; training rate; lowest of the training and deployment rates |
| $s$ | output stride (Section 4.6) |
| $B$; $W_0=f_{\min}-B$ | signal band; the critical frequency above which kernel energy aliases at some admissible rate |
| $D$, $T=D/2$, $I=[-T,T]$ | kernel support |
| $W$, $P$ | prolate band; number of basis functions |
| $\varphi_p$, $\lambda_p$ | prolate spheroidal wave functions and their concentration eigenvalues |
| $a_p$ | coefficients of a kernel in the prolate basis |
| $k$, $K$; $h_f$, $H_f$ | continuous kernel and its spectrum; taps and discrete response at rate $f$ |
| $m$ | alias order |
| $\mathcal{E}_\Omega(k)$ | kernel energy on the frequency set Ω |
| $\mathrm{Err}_f$, $\overline{\mathrm{Err}}_f$ | fixed-grid and offset-averaged deployment error |
| $c(k)$, $c(V)$ | transfer certificate of a kernel; worst-case certificate of a space |
| $\gamma_f$, $\Gamma_f$ | fixed-grid constants of a space |
| $S$ | input power spectrum used as a weight |
| $j$ | filter index (Section 4) |

**Kernels.** A temporal kernel is a function $k\colon \mathbb{R}\to\mathbb{C}$ with $\operatorname{supp} k \subseteq I := [-T,T]$ and $T = D/2$, where $D = 0.25$ s. The results that sample $k$ at points, namely Theorem S3.1, Proposition S5.1 and Theorems S8.1 and S9.1, assume that $k$ has bounded variation on $\mathbb{R}$ and that its zero extension takes midpoint values, $k(t) = 1/2\,\{k(t^+) + k(t^-)\}$ for every $t$. Theorem S4.1, by contrast, needs only $k \in$

$L^2(\mathbb{R})$. We write $k_I$ for the restriction of $k$ to $(-T, T)$, extended to $\pm T$ by its one-sided limits. For a PKC kernel, $k_I$ is the PSWF expansion on the closed interval, and the midpoint normalization changes only the end values, which become $k(\pm T) = 1/2\, k_I(\pm T)$.

**Spectra and band energies.** We write $K(\xi) = \int_{\mathbb{R}} k\,(t) e^{-2\pi i \xi t} dt$. Because $k$ is square integrable with bounded support, $K$ extends to an entire function, and Plancherel's theorem gives $\|K\|_2 = \|k\|_2$. For a measurable set $\Omega \subseteq \mathbb{R}$ we write $\mathcal{E}_\Omega(k) = \int_\Omega |K|^2$, together with $\mathcal{E}_{\geq V}(k) = \mathcal{E}_{\{|\xi| \geq V\}}(k)$, $\mathcal{E}_{>V}(k) = \mathcal{E}_{\{|\xi|>V\}}(k)$ and $\mathcal{E}_{[a,b]}(k) = \mathcal{E}_{\{a \leq |\xi| \leq b\}}(k)$.

**Realization at rate $f$.** The taps are $h_f[n] = k(n/f)/f$ for $n \in \mathbb{Z}$. Under the midpoint normalization this is the implemented rule $h_f[n] = w_n k_I(n/f)/f$ at the exact physical lags $n/f \in I$, with the trapezoid end weight $w_n = 1/2$ when $n/f = \pm T$ and $w_n = 1$ otherwise. The implementation multiplies each kernel by a gain that is computed once at the training rate and kept at every rate, and we absorb this constant into $k$. The implementation also applies the taps as a cross-correlation, which realizes the reflected kernel $k(-t)$. All frequency sets below are symmetric and all weights are even, so every quantity takes the same value for $k$ and for its reflection. The PKC spaces are themselves invariant under reflection, because each $\varphi_p$ is even or odd. The realized response is the $f$-periodic trigonometric polynomial $H_f(\nu) = \sum_n h_f\,[n] e^{-2\pi i \nu n/f}$, and we write $e_f = H_f - K$.

**Data band and rates (assumption A1).** The input of the layer is band-limited to $[-B, B]$, and every rate satisfies $f \geq f_{\min} > 2B$. We write $W_0 := f_{\min} - B$, where $f_{\min}$ is the lowest of the training and deployment rates. In this paper $B = 45$ Hz, $f_{\min} = 100$ Hz and $W_0 = 55$ Hz. For an input whose spectrum extends to $B'$ with $B < B' < f_{\min}/2$, every statement below holds with $B$ replaced by $B'$ and $W_0$ by $f_{\min} - B'$.

**Lemma S2.1 (what a realized layer computes) (classical).** Assume A1. Let $x(t) = \int_{-B}^{B} X\,(\nu) e^{2\pi i \nu t} d\nu$ with $X \in L^2(-B, B)$, and let $y[q] = \sum_n h_f\,[n] x((q - n)/f)$. Then $y[q] = y_f(q/f)$ with $y_f(t) = \int_{-B}^{B} X\,(\nu) H_f(\nu) e^{2\pi i \nu t} d\nu$. The continuous layer outputs $y = k * x$ with $\hat{y} = XK$, and the two sampled outputs satisfy

$$\frac{1}{f}\sum_{q\in\mathbb{Z}} | y_f(q/f) - y(q/f)|^2 = \int_{-B}^{B} | X(\nu)|^2 |e_f(\nu)|^2 d\nu.$$

*Source.* The lemma combines the sampling theorem (Shannon, 1949) with Parseval's identity.

*Proof.* Inserting the Fourier representation of $x$ into the finite sum, we obtain $y[q] = \int X(\nu) \sum_n h_f [n] e^{2\pi i\nu(q-n)/f} d\nu = y_f(q/f)$. The function $g = y_f - y$ has the spectrum $X(H_f - K)$, supported in $[-B,B] \subset (-f/2, f/2)$. Hence $g(q/f) = \int_{-f/2}^{f/2} \hat{g}\,(\nu) e^{2\pi i\nu q/f} d\nu$, and Parseval's identity for the orthonormal basis $\{f^{-1/2} e^{2\pi i\nu q/f}\}_{q\in\mathbb{Z}}$ of $L^2(-f/2, f/2)$ yields $\sum_q | g(q/f)|^2 = f \|\hat{g}\|_2^2$. □

**Deployment error.** Let $S \geq 0$ be an even, bounded and measurable weight on $[-B,B]$, extended by zero, which stands for the input power spectrum ($S \equiv 1$ when no spectrum is used). We define

$$\mathrm{Err}_f^S(k) = \int_{-B}^{B} S\,(\nu) |e_f(\nu)|^2 d\nu,$$
$$\mathrm{Err}_f = \mathrm{Err}_f^1.$$

The error functional Err is unrelated to the support length $D$. By Lemma S2.1, an input with $|X|^2 = \sigma^2 S$ produces the output error energy $\sigma^2 \mathrm{Err}_f^S(k)$. Translating the input multiplies $X$ by a unimodular factor, so this energy does not depend on where the input samples fall. Between a training rate $f_0$ and a deployment rate $f$ the realized response changes by $H_f - H_{f_0} = e_f - e_{f_0}$.

**Alias bands and guard band.** We set $A_f = \bigcup_{m\neq 0}(mf + [-B,B])$ and, for a set of rates $F$, $\Omega_F = \bigcup_{f\in F} A_f$. For $m \neq 0$ and $|\nu| \leq B$ we have $|mf + \nu| \geq f - B$, hence

$$A_f \subseteq \{|\xi| \geq f - B\} \subseteq \{|\xi| \geq W_0\} \quad (f \geq f_{\min}). \tag{S1}$$

The edge $f - B$ is the stopband edge that multistage decimators use so that no component folds into the baseband (Crochiere & Rabiner, 1975). The guard band $\{B < |\xi| < W_0\}$ lies outside the data band and meets no alias band $A_f$ with $f \geq f_{\min}$, so kernel energy in the guard band enters no deployment error at any admissible rate.

**Concentration operator and prolate functions.** For $V > 0$ let $(Q_V k)(t) = \int_{-T}^{T} \frac{\sin 2\pi V(t-s)}{\pi(t-s)} k(s) ds$ on $L^2(I)$. Plancherel's theorem gives $\langle Q_V k, k\rangle = \int_{-V}^{V} |K|^2$, so that

$$\mathcal{E}_{\geq V}(k) = ||k||^2 - \langle Q_V k, k\rangle \quad (k \in L^2(I)). \tag{F1}$$

The operator $Q_V$ is compact, self-adjoint and positive. Its eigenvalues $1 > \lambda_1(D,V) > \lambda_2(D,V) > \cdots > 0$ are simple and belong to orthonormal eigenfunctions $\varphi_p(\cdot; D, V)$, the time-limited prolate spheroidal wave functions (PSWFs) of Slepian and Pollak (1961). We index from 1, whereas Slepian and Pollak index from 0. We use the counting function $N_\alpha(D,V) = \#\{p: \lambda_p(D,V) \geq \alpha\}$ and the spaces $\mathcal{V}_P(D,V) = \mathrm{span}\{\varphi_1, \dots, \varphi_P\}$. For $k = \sum_p a_p\, \varphi_p(\cdot; D, V)$, (F1) and orthonormality give

$$\mathcal{E}_{\geq V}(k) = \textstyle\sum_p \big(1 - \lambda_p(D,V)\big)\, |a_p|^2. \tag{F1′}$$

A prolate kernel convolution (PKC) kernel is $k = \sum_{p \leq P} a_p\, \varphi_p(\cdot; D, W)$. The main design uses $W = W_0 = 55$ Hz and $P = 25 = N_{0.99}(D, 55)$.

(F2) If $0 \neq k \in L^2(I)$, then $K$ is a nonzero entire function, whose zeros are isolated, so $\mathcal{E}_\Omega(k) > 0$ for every $\Omega$ of positive measure.

**Certificates.** For a kernel $k \neq 0$ the transfer certificate of Section 4.5 is $c(k) = \mathcal{E}_{\geq W_0}(k)/||k||^2$. For a finite-dimensional subspace $\mathcal{V} \subset L^2(I)$ we define the worst-case certificate

$$c(\mathcal{V}) = \max_{0 \neq k \in \mathcal{V}} c(k) = \max_{0 \neq k \in \mathcal{V}} \frac{\mathcal{E}_{\geq W_0}(k)}{||k||^2}.$$

By (S1) and Lemma A.1, $c(\mathcal{V})$ is the largest fraction of kernel energy that a member of $\mathcal{V}$ places in the frequencies that alias into the data band at some rate of a covering rate set. By (F2), $c(\mathcal{V}) > 0$. For the main PKC space, $c = 1 - \lambda_{25}(D, 55) = 9.381 \times 10^{-3}$ (Proposition 2(i)). Now let the input spectrum $S$ not vanish almost everywhere on $[-B, B]$. Then, for a single kernel $k \neq 0$, the denominator below is positive by (F2), and we define the relative certificate

$$c_S(k) = \frac{||S||_\infty \mathcal{E}_{\geq W_0}(k)}{\int_{-B}^{B} S\, |K|^2}.$$

We compute it from the trained weights and the training spectrum $S$. It bounds the offset-averaged error, relative to the in-band output energy of the continuous layer, at every rate $f \geq f_{\min}$ (Theorem S4.1(*f*)).

### S3. The rate-transfer identity

**Theorem S3.1 (rate-transfer identity) (classical).** Let $k$ have bounded variation and bounded support, normalized to midpoint values. For every $f > 0$ and every $\nu \in \mathbb{R}$,

$$H_f(\nu) = \lim_{M\to\infty} \sum_{m=-M}^{M} K(\nu + mf).$$

Consequently, for $|\nu| \le B$, the error $e_f(\nu) = \lim_{M\to\infty} \sum_{0<|m|\le M} K(\nu + mf)$ depends on $K$ only through its values on $A_f$, and $H_f - H_{f_0} = e_f - e_{f_0}$.

*Source.* Theorem S3.1 is the Poisson summation formula (Poisson, 1823, 1827) for functions of bounded variation with bounded support. It follows from the Dirichlet–Jordan theorem: the Fourier series of a periodic function of bounded variation converges at every point to the mean of the one-sided limits. Dirichlet (1829) proved this for piecewise monotone functions and Jordan (1881) for functions of bounded variation, and Zygmund (2003, Vol. I) gives a modern statement.

*Proof.* We fix $f$ and $\nu$ and put $\varphi(t) = k(t)e^{-2\pi i\nu t}$, a function of bounded variation with bounded support. Its periodization $\Phi(t) = \sum_{n\in\mathbb{Z}} \varphi(t + n/f)$ is a finite sum on bounded sets, has period $1/f$ and has bounded variation on a period. Since the translation $t \mapsto t + n/f$ leaves $e^{-2\pi imft}$ unchanged, the Fourier coefficients of $\Phi$ are

$$\widehat{\Phi}_m = f\int_0^{1/f} \Phi(t)e^{-2\pi imft}dt = f\int_{\mathbb{R}} \varphi(t)e^{-2\pi imft}dt = fK(\nu + mf).$$

By the Dirichlet–Jordan theorem, $\sum_{|m|\le M} \widehat{\Phi}_m e^{2\pi imft} \to \frac{1}{2}\{\Phi(t^+) + \Phi(t^-)\}$ at every $t$. At $t = 0$ the limit equals $\sum_n \frac{1}{2}\{\varphi(n/f^+) + \varphi(n/f^-)\} = \sum_n k(n/f)e^{-2\pi i\nu n/f} = fH_f(\nu)$, by the midpoint normalization and the continuity of $e^{-2\pi i\nu t}$. Dividing by $f$, we obtain the identity. For $|\nu| \le B$ the term $m = 0$ is $K(\nu)$, and $\nu + mf \in A_f$ for $m \ne 0$. □

**Remark S3.1 (end weight and convergence).** (i) The midpoint normalization is the trapezoid end weight, the same half weight that corrects impulse-invariant filter design when the impulse response jumps at the origin (Jackson, 2000; Mecklenbräuker, 2000). With full weights on the end taps, the realized response becomes $H_f(\nu) + \frac{1}{2f}\{k_I(T)e^{-2\pi i\nu T} + k_I(-T)e^{2\pi i\nu T}\}$ whenever $Tf \in \mathbb{Z}$, which holds at 128, 160, 200 and 256 Hz. This end-point term does not arise from aliasing, and it decays like $1/f$. The tap set changes when $Tf$ crosses an integer, so $H_f$, as a function of the rate, jumps at every $f$ with $Tf \in \mathbb{Z}$ ($f \in 8\mathbb{Z}$ Hz for $D = 0.25$ s) unless $k_I(T) = k_I(-T) = 0$. At such a rate its one-sided limits are $H_f$ minus and plus the end-point term: from below the end taps are absent, and from above they carry full weight. The fixed-grid error $\mathrm{Err}_f$ therefore jumps at these rates (Proposition S5.1(b)), whereas the offset average $\overline{\mathrm{Err}}_f = \mathcal{E}_{A_f}(k)$ of Theorem S4.1(c) depends continuously on $f$ by dominated convergence. (ii) PSWFs do not vanish at $\pm T$, so a PKC kernel has $|K(\xi)|$ of order $|\xi|^{-1}$, and Theorem S3.1 needs the symmetric partial sums. When the zero extension of $k$ belongs to $H^1(\mathbb{R})$, the series converges absolutely at almost every $\nu$ (proof of Proposition S5.1(c)).

*Numerical check.* We compared $H_f$ with the symmetric partial sums for a random unit-norm kernel of the main PKC space. At $|m| \le 2000$ the relative difference lies between $2.0 \times 10^{-11}$ at 500 Hz and $1.2 \times 10^{-6}$ at 128 Hz. From $M = 100$ to $M = 2000$ it falls by a factor of at least 20, and a piecewise-linear kernel with end jumps gives at most $6.0 \times 10^{-5}$. With full end weights, the discrepancy at the four rates with $Tf \in \mathbb{Z}$ equals the end-point term of (i), for example $8.52 \times 10^{-4}$ at 128 Hz. At the other four rates it stays below $7 \times 10^{-8}$ for the PKC kernel. We also checked the one-sided limits of $e_f$ at 104 Hz for a random unit kernel of the main space. They agree with $e_{104}$ minus and plus the end-point term to within $2.5 \times 10^{-10}$, whereas the end-point term itself reaches magnitudes up to $9.7 \times 10^{-4}$.

### S4. Offset-averaged deployment error

**Definition S4.1 (offset realization and offset average).** Let $k \in L^2(\mathbb{R})$ vanish outside a bounded interval, and fix a representative of $k$ (the midpoint-normalized one when $k$ has bounded variation). For $\theta \in [0,1)$, the $\theta$-offset realization at rate $f$ has the taps $h_{f,\theta}[n] = k((n+\theta)/f)/f$, which act at the physical lags $t_n = (n+\theta)/f$ . Its response referred to physical time is $H_{f,\theta}(\nu) = \sum_n h_{f,\theta}\,[n] e^{-2\pi i \nu t_n}$, and $e_{f,\theta} = H_{f,\theta} - K$. The implemented layer is the realization with $\theta = 0$. With $\mathrm{Err}^S_{f,\theta}(k) = \int_{-B}^{B} S\,|e_{f,\theta}|^2$, the offset-averaged deployment error is

$$\overline{\mathrm{Err}}_f^S(k) = \int_0^1 \mathrm{Err}_{f,\theta}^S(k)d\theta,$$

$$\overline{\mathrm{Err}}_f = \overline{\mathrm{Err}}_f^1.$$

Two representatives of $k$ give the same taps for almost every $\theta$, so $\overline{\mathrm{Err}}_f^S$ depends on $k$ only as an element of $L^2$.

**Lemma S4.1 (the offset average on a deployment grid) (classical).** Let $k$ have bounded variation and midpoint values, let $k_{t_0} = k(\cdot + t_0)$ for $t_0 \in \mathbb{R}$, and write $ft_0 = q + \theta$ with $q \in \mathbb{Z}$ and $\theta \in [0,1)$.

(i) The implemented realization of $k_{t_0}$ at rate $f$ has the taps $h_f[k_{t_0}][n] = h_{f,\theta}[n+q]$ and the error $e_f[k_{t_0}](\nu) = e^{2\pi i\nu t_0} e_{f,\theta}(\nu)$, so that $\mathrm{Err}_f^S(k_{t_0}) = \mathrm{Err}_{f,\theta}^S(k)$.

(ii) The function $t_0 \mapsto \mathrm{Err}_f^S(k_{t_0})$ has period $1/f$, and

$$\overline{\mathrm{Err}}_f^S(k) = f\int_0^{1/f} \mathrm{Err}_f^S(k_{t_0})dt_0$$
$$= \lim_{L\to\infty} \frac{1}{2L}\int_{-L}^{L} \mathrm{Err}_f^S(k_{t_0})dt_0.$$

(iii) For every input $x$ and every $t_1 \in \mathbb{R}$, the inputs $x$ and $x(\cdot - t_1)$ have the same output error energy in Lemma S2.1.

On a deployment grid of rate $f$, the offset average is the implemented-grid error averaged uniformly over all placements of the kernel relative to the lag lattice $f^{-1}\mathbb{Z}$. The implemented layer is the placement $t_0 = 0$, which puts the kernel centre on a lag, whereas the phase of the input samples plays no role.

*Source.* The lemma is a change of variables: a uniformly random origin of the sampling lattice is equivalent to a uniformly random translation of the integrand relative to a fixed lattice. This equivalence is the set-up of systematic sampling with a random origin (Moran, 1950).

*Proof.* (i) We have $k_{t_0}(n/f) = k((n+q+\theta)/f) = fh_{f,\theta}[n+q]$. Hence

$$H_f[k_{t_0}](\nu)$$
$$= \sum_n h_{f,\theta}[n+q]e^{-2\pi i\nu n/f}$$
$$= e^{2\pi i\nu(q+\theta)/f}\sum_{n'} h_{f,\theta}[n']e^{-2\pi i\nu(n'+\theta)/f} = e^{2\pi i\nu t_0}H_{f,\theta}(\nu)$$

The continuous response of $k_{t_0}$ is $e^{2\pi i\nu t_0}K(\nu)$. (ii) Replacing $t_0$ by $t_0+1/f$ replaces $q$ by $q+1$ and keeps $\theta$, so the function has period $1/f$ by (i). On $[0,1/f)$ we have $q=0$ and $\theta=ft_0$, and we substitute $\theta=ft_0$ to obtain the first equality; periodicity gives the second. (iii) Translation multiplies $X$ by $e^{-2\pi i\nu t_1}$, which leaves $|X|^2$, and hence the right-hand side of Lemma S2.1, unchanged. □

**Theorem S4.1 (offset-averaged deployment error) (classical: (a), (b), (d), and (c) for $S\equiv 1$; new: (c) with a spectrum weight, (e)–(g)).** Let $k\in L^2(\mathbb{R})$ vanish outside a bounded interval, let $f>0$, and let $S$ be a weight as above.

(a) For every $\nu\in\mathbb{R}$ the function $\theta\mapsto H_{f,\theta}(\nu)$ belongs to $L^2(0,1)$, and its Fourier coefficients are

$$\int_0^1 H_{f,\theta}(\nu)e^{-2\pi i m\theta}d\theta = K(\nu+mf)$$
$$(m\in\mathbb{Z}).$$

If $k$ has bounded variation and midpoint values, then $e_{f,\theta}(\nu)=\lim_{M\to\infty}\sum_{0<|m|\le M}K(\nu+mf)e^{2\pi i m\theta}$ for all $\theta$ and $\nu$.

(b) For every $\nu$, $\int_0^1|e_{f,\theta}(\nu)|^2d\theta=\sum_{m\neq 0}|K(\nu+mf)|^2$ and $\int_0^1 e_{f,\theta}(\nu)d\theta=0$.

(c) (Symbol form.) $\overline{\mathrm{Err}}_f^S(k)=\int_{\mathbb{R}} w_f^S(\xi)|K(\xi)|^2d\xi$ with $w_f^S(\xi)=\sum_{m\neq 0}S(\xi-mf)$. If $f\ge 2B$, then $0\le w_f^S\le\|S\|_\infty\mathbf{1}_{A_f}$ almost everywhere, and $\overline{\mathrm{Err}}_f(k)=\mathcal{E}_{A_f}(k)$: for a flat input spectrum the offset-averaged deployment error equals the alias-band energy.

(d) (Lattice form.) Let $\hat{s}(\tau)=\int S(\nu)e^{-2\pi i\nu\tau}d\nu$, and let $r_k(\tau)=\int k(t+\tau)\overline{k(t)}dt$ be the autocorrelation of $k$. Then

$$\overline{\mathrm{Err}}_f^S(k)=\frac{1}{f}\sum_{p\in\mathbb{Z}}\hat{s}(p/f)r_k(p/f)-\int_{\mathbb{R}}\hat{s}(\tau)r_k(\tau)d\tau,$$

which is the error of the rectangle rule with step $1/f$ for the integral of $\hat{s}r_k$. When $\mathrm{supp}k\subseteq I$, only the terms with $|p|<Df$ are nonzero. For $S\equiv 1$, $\hat{s}(\tau)=2B\mathrm{sinc}(2B\tau)$ with $\mathrm{sinc}x=\sin(\pi x)/(\pi x)$.

(e) (Two rates.) For $f,f_0>0$ and offsets $\theta,\theta'$ that are independent and uniform on $[0,1)$,

$$\int_0^1\int_0^1\int_{-B}^{B}S\,|H_{f,\theta}-H_{f_0,\theta'}|^2d\nu d\theta d\theta'=\overline{\mathrm{Err}}_f^S(k)+\overline{\mathrm{Err}}_{f_0}^S(k).$$

(*f*) (Bound by the energy beyond $f_{\min}-B$.) If $f\ge f_{\min}$, then

$$\overline{\mathrm{Err}}_f^S(k) \le \|S\|_\infty \mathcal{E}_{A_f}(k) \le \|S\|_\infty \mathcal{E}_{\ge f-B}(k) \le \|S\|_\infty \mathcal{E}_{\ge W_0}(k).$$

Hence $\overline{\mathrm{Err}}_f^S(k) \le \|S\|_\infty c(\mathcal{V})\|k\|^2$ for every member $k$ of a space $\mathcal{V}$, $\overline{\mathrm{Err}}_f^S(k) \le c_S(k)\int S|K|^2$, and the two-rate average of (e) is at most $2\|S\|_\infty \mathcal{E}_{\ge W_0}(k)$ when $f, f_0 \ge f_{\min}$.

(g) (Attainment.) $\overline{\mathrm{Err}}_{f_{\min}}(k) \ge \mathcal{E}_{\ge W_0}(k) - \mathcal{E}_{>f_{\min}+B}(k)$.

*Proof.* (a) Only finitely many $n$ contribute to $H_{f,\theta}(\nu)$, and for each of them the substitution $t = (n+\theta)/f$ gives $\int_0^1 |k((n+\theta)/f)|^2 d\theta = f\int_{n/f}^{(n+1)/f} |k|^2 < \infty$, so $\theta \mapsto H_{f,\theta}(\nu)$ lies in $L^2(0,1)$. The same substitution, with $e^{-2\pi i m\theta} = e^{-2\pi i m f t}$ because $e^{2\pi i m n} = 1$, yields

$$\int_0^1 H_{f,\theta}(\nu) e^{-2\pi i m\theta} d\theta = \sum_n \int_{n/f}^{(n+1)/f} k(t) e^{-2\pi i(\nu+mf)t} dt = K(\nu + mf).$$

For the pointwise statement, the kernel $k_{\theta/f} = k(\cdot + \theta/f)$ has bounded variation, bounded support, midpoint values and spectrum $e^{2\pi i\xi\theta/f}K(\xi)$. By Lemma S4.1(i) its implemented response is $e^{2\pi i\nu\theta/f}H_{f,\theta}(\nu)$, and Theorem S3.1 gives $H_{f,\theta}(\nu) = \lim_M \sum_{|m|\le M} e^{2\pi i m\theta} K(\nu + mf)$, whose term $m = 0$ is $K(\nu)$.

(b) By (a), the function $\theta \mapsto e_{f,\theta}(\nu) = H_{f,\theta}(\nu) - K(\nu)$ has the Fourier coefficients $K(\nu + mf)$ for $m \ne 0$ and 0 for $m = 0$. Parseval's identity in $L^2(0,1)$ gives both statements.

(c) The integrand $S(\nu)|e_{f,\theta}(\nu)|^2 \ge 0$ is jointly measurable in $(\theta, \nu)$, so we apply Tonelli's theorem and (b) and obtain $\overline{\mathrm{Err}}_f^S(k) = \sum_{m\ne 0}\int S(\nu)|K(\nu + mf)|^2 d\nu = \sum_{m\ne 0}\int S(\xi - mf)|K(\xi)|^2 d\xi$, which is the symbol form. If $f \ge 2B$, the intervals $mf + [-B, B]$ with $m \ne 0$ overlap at most at end points. Almost every $\xi$ then meets at most one nonzero term of $w_f^S$, and $w_f^S$ vanishes off $A_f$. For $S \equiv 1$ this gives $w_f^1 = \mathbf{1}_{A_f}$ almost everywhere.

(d) For $m \in \mathbb{Z}$ let $G_m = \int S(\xi - mf)|K(\xi)|^2 d\xi \ge 0$. Since $|K(\xi)|^2 = \int r_k(\tau) e^{-2\pi i\xi\tau} d\tau$ and $\int S(\xi - mf) e^{-2\pi i\xi\tau} d\xi = e^{-2\pi i m f\tau}\hat{s}(\tau)$, Fubini's theorem gives $G_m = \int g_S(\tau) e^{-2\pi i m f\tau} d\tau$ with $g_S = \hat{s} r_k$. The function $g_S$ is continuous with bounded support, so its $1/f$-periodization $\sum_p g_S(\tau + p/f)$ is continuous, with Fourier coefficients $fG_m$. These coefficients are summable, because

$\sum_m G_m = \int (\sum_m S(\xi - mf))|K|^2 \le (2B/f + 1)\|S\|_\infty \|k\|^2$. A continuous periodic function with summable Fourier coefficients equals the sum of its Fourier series at every point, and at $\tau = 0$ this reads $\sum_p g_S(p/f) = f \sum_m G_m$. We subtract $G_0 = \int S|K|^2 = \int g_S$ and use (c) to obtain the lattice form. When $\mathrm{supp} k \subseteq I$, $r_k$ vanishes for $|\tau| \ge D$.

(e) We fix $\nu$ and write $u(\theta) = e_{f,\theta}(\nu)$ and $v(\theta') = e_{f_0,\theta'}(\nu)$, so that $H_{f,\theta} - H_{f_0,\theta'} = u - v$. Both functions have zero mean by (b), hence $\iint |u - v|^2 = \int |u|^2 + \int |v|^2$. Integrating against $S$ and applying Tonelli's theorem completes the proof.

(*f*) Since $f \ge f_{\min} > 2B$, (c) gives the first inequality, (S1) the next two, and the definitions of $c$ and $c_S$ the remaining bounds. The two-rate bound follows from (e).

(g) By (c), $\overline{\mathrm{Err}}_{f_{\min}}(k) = \mathcal{E}_{A_{f_{\min}}}(k)$, and the bands $m = \pm 1$ of $A_{f_{\min}}$ cover $W_0 \le |\xi| \le f_{\min} + B$. □

*Sources and prior work.* Parts (a) and (b) are the classical analysis of systematic sampling with a uniformly random origin, applied to the integrand $t \mapsto k(t)e^{-2\pi i \nu t}$. Moran (1950) showed that the shifted rectangle sum over $\mathbb{R}$ is unbiased and evaluated its variance as a function of the step. This variance equals the sum of the squared nonzero Fourier coefficients of the periodized integrand, the one-dimensional case of the variance formula for randomly shifted lattice rules (L'Ecuyer & Lemieux, 2000). Such random shifts go back to Cranley and Patterson (1976). Part (d) is Matheron's transitive formula, which writes the variance of the same estimator as the rectangle-rule error of the covariogram of the integrand (Matheron, 1970/2019; Cruz-Orive, 1989). Here the covariogram is $e^{-2\pi i \nu \tau} r_k(\tau)$, integrated against $S$. Part (c) with $S \equiv 1$ is the shift-averaged error formula of Blu and Unser (1999, Theorem 2) for sampling without prefilter, restricted to the data band. Earlier, Park and Schowengerdt (1982) separated the phase-averaged aliasing of sampled imaging systems from its variation over the phase in the same way. Brown (1978) bounded the mean-square aliasing error of the cardinal series of a wide-sense stationary process by a constant times its spectral power outside the band, uniformly in time, and showed that this constant is the best possible. For a stationary process the alias orders are uncorrelated and add in power. A learned kernel, however, is deterministic, and at a fixed offset its alias orders $K(\nu + mf)$ add in amplitude; averaging over the offset makes them orthogonal through the characters $e^{2\pi i m\theta}$. Beyond these classical results, Theorem S4.1 contributes for deployment the spectrum-weighted symbol on the data band, the two-rate identity (e) and the rate-uniform bounds (*f*) and (g) by the energy beyond $f_{\min} - B$. These bounds rest on the offset average, since at a fixed offset no bound with a universal constant holds (Proposition S5.1(a)).

*Numerical check.* With 2048 offsets and $S \equiv 1$, the offset average and the alias-band energy agree to within a relative error of $5.2 \times 10^{-6}$ at all eight rates for two PKC kernels, one with $W = 55$ Hz and $P = 25$ and one with $W = 45$ Hz and $P = 31$. The agreement is $2.4 \times 10^{-5}$ for a piecewise-linear kernel with end jumps and $5.9 \times 10^{-10}$ for a smooth bump. The offset mean of $e_{f,\theta}$ stays below $2.2 \times 10^{-7}$ of its root-mean-square value. We repeated the comparison with 1024 offsets and the model spectrum $S(\nu) = 1/(1 + (\nu/8)^2)$. The offset average, the symbol form and the lattice form then agree to $5.1 \times 10^{-6}$ for the main PKC kernel and to $1.2 \times 10^{-5}$ for the kernel with $W = 45$ Hz and $P = 31$. For the bump they agree to $5.7 \times 10^{-9}$, again at all eight rates. For a random unit kernel of the main space, $\overline{\mathrm{Err}}_{100} = 9.197 \times 10^{-5}$ stays below the kernel's energy beyond 55 Hz, which is $9.402 \times 10^{-5}$.

**Corollary S4.1 (new).**

(a) (Main space.) Every kernel of $\mathcal{V}_{25}(D, 55)$ satisfies $\overline{\mathrm{Err}}_f^S(k) \leq c\|S\|_\infty \|k\|^2$ with $c = 9.381 \times 10^{-3}$ at every rate $f \geq 100$ Hz, and its two-rate average in Theorem S4.1(e) is at most $2c\|S\|_\infty \|k\|^2 < 1.88 \times 10^{-2} \|S\|_\infty \|k\|^2$. At 100 Hz the largest alias-band energy over unit kernels of this space equals $0.9785c$, so the certificate is attained there to within 2.2%.

(b) (Energy just above $W_0$ reaches only the top of the data band.) Let $0 < \delta < f_{\min}$, let $f \geq f_{\min}$, and let $s_\delta = \operatorname{esssup}\{S(\nu): B - \delta \leq |\nu| \leq B\}$. Then $w_f^S(\xi) \leq s_\delta$ for $W_0 \leq |\xi| \leq W_0 + \delta$, and

$$\overline{\mathrm{Err}}_f^S(k) \leq s_\delta \mathcal{E}_{[W_0, W_0+\delta]}(k) + \|S\|_\infty \mathcal{E}_{>W_0+\delta}(k).$$

*Proof.* (a) This follows from Theorem S4.1(e) and (*f*) with $c = c(\mathcal{V}_{25}(D, 55)) = 1 - \lambda_{25}(D, 55)$. (b) Let $W_0 \leq \xi \leq W_0 + \delta$; negative $\xi$ follow by symmetry. For $m \geq 2$ we have $\xi - mf \leq W_0 + \delta - 2f_{\min} = \delta - f_{\min} - B < -B$, and for $m \leq -1$ we have $\xi - mf \geq W_0 + f_{\min} > B$. Only $m = 1$ contributes, with $\nu = \xi -$

$f \le W_0 + \delta - f_{\min} = \delta - B$. Whenever $S(\nu) \neq 0$ we also have $\nu \ge -B$, so $B - \delta \le |\nu| \le B$ and $w_f^S(\xi) = S(\xi - f) \le s_\delta$. Below $W_0$ the symbol vanishes by (S1), and $w_f^S \le \|S\|_\infty$ everywhere because $f \ge 2B$. Inserting these bounds into the symbol form, we obtain the inequality. □

Take the model spectrum $S(\nu) = 1/(1 + (\nu/8)^2)$ and $\delta = 10$ Hz. Then $s_\delta = S(35) = 0.050$, so kernel energy in 55–65 Hz enters the offset-averaged error with weight at most 0.050, whereas energy near a multiple of the rate enters with weight close to 1. A kernel whose energy beyond $W_0$ sits just above $W_0$ thus transfers with an error far below its flat certificate when the input spectrum is weak at the top of the data band.

### S5. The implemented grid

**Proposition S5.1 (fixed grid).**

(a) **No universal constant.** Let $\phi(x) = \sqrt{8/3}\cos^2(\pi x)$ for $|x| \le \frac{1}{2}$ and $\phi(x) = 0$ otherwise. Then $\phi \in C^1$, $\|\phi\|_2 = 1$, $\|\phi\|_1 = \sqrt{2/3}$ and $\phi(0) = 2\|\phi\|_1$. For $f \ge 2B$ and $0 < \delta < \min\{2/f, D\}$ let $k_\delta(t) = \delta^{-1/2}\phi(t/\delta)$. Then $k_\delta \in C^1$, $k_\delta(\pm T) = 0$, $\|k_\delta\| = 1$ and $\overline{\mathrm{Err}}_f(k_\delta) = \mathcal{E}_{A_f}(k_\delta) \le 1$, whereas

$$\mathrm{Err}_f(k_\delta) \ge \frac{4B}{3}\frac{(2 - f\delta)^2}{f^2\delta} \to \infty$$
$$(\delta \to 0).$$

The failure persists under a constraint on the out-of-band energy. Let $\eta > 1 - \lambda_1(D, W_0)$; for our $D$ and $W_0$, $1 - \lambda_1$ lies below $1 - \lambda_{15}(D, 55) = 6.3 \times 10^{-13}$. Then there are $C^1$ kernels $u_\delta$ with $u_\delta(\pm T) = 0$, $\|u_\delta\| \le 2$ and $\mathcal{E}_{\ge W_0}(u_\delta) \le \eta\|u_\delta\|^2$ for all small $\delta$, whose fixed-grid error $\mathrm{Err}_f(u_\delta)$ tends to infinity. Hence no inequality $\mathrm{Err}_f(k) \le C\mathcal{E}_{A_f}(k)$ or $\mathrm{Err}_f(k) \le C\mathcal{E}_{\ge W_0}(k)$ with a constant $C$ independent of $k$ holds, and a small out-of-band energy does not bound the error at a fixed grid, even for $C^1$ kernels that vanish at $\pm D/2$.

*Proof.* The support $[-\delta/2, \delta/2]$ lies in $(-1/f, 1/f)$, so the only lag in the support is $n = 0$, and $H_f(\nu) = k_\delta(0)/f = \phi(0)/(f\sqrt{\delta})$ for all $\nu$. Further, $|K_\delta(\nu)| \le \|k_\delta\|_1 = \sqrt{\delta}\|\phi\|_1$. Hence $|e_f(\nu)| \ge \|\phi\|_1(2 - f\delta)/(f\sqrt{\delta}) > 0$ on $[-B, B]$, and we integrate the square over $[-B, B]$ with $\|\phi\|_1^2 = 2/3$ to obtain the lower bound. The energy statements follow from Plancherel's theorem and Theorem S4.1(c). For the second part, $\mathcal{E}_{\ge W_0}(\varphi_1) = 1 - \lambda_1(D, W_0) < \eta$, and $C^\infty$

functions supported in $(-T,T)$ are dense in $L^2(I)$, so there is a unit $C^\infty$ kernel $u$ supported in $(-T,T)$ with $\mathcal{E}_{\geq W_0}(u) < \eta$. We set $u_\delta = u + \delta^{1/4} k_\delta$. Then $\|u_\delta\| \to 1$ and $\mathcal{E}_{\geq W_0}(u_\delta) \to \mathcal{E}_{\geq W_0}(u)$, so $\mathcal{E}_{\geq W_0}(u_\delta) \leq \eta \|u_\delta\|^2$ for small $\delta$. Since $e_f$ is linear in the kernel, $\mathrm{Err}_f(u_\delta)^{1/2} \geq \delta^{1/4}\mathrm{Err}_f(k_\delta)^{1/2} - \mathrm{Err}_f(u)^{1/2}$, which grows like $\delta^{-1/4}$. □

*Numerical check.* At $f = 100$ Hz we decreased $\delta$ from $8 \times 10^{-3}$ to $2.5 \times 10^{-4}$. Over this range the ratio $\mathrm{Err}_{100}/\mathcal{E}_{A_{100}}$ grows from 2.5 to 106 in proportion to $1/\delta$, while $\mathcal{E}_{A_{100}}$ rises only from 0.45 to 0.89. The computed fixed-grid error lies within 4% of the lower bound at every $\delta$.

*Prior work.* For inputs band-limited to the span of a multiband spectral support, Venkataramani and Bresler (2000) bound the peak value and the energy of the aliasing error. A time-limited kernel, however, is band-limited to no finite interval, and part (a) shows that its fixed-grid error admits no bound by a universal constant times its out-of-band energy. Part (b) recovers finite constants by restricting the kernel to a finite-dimensional space.

(b) **Space constants.** Let $\mathcal{V} \subset L^2(I)$ be a nonzero finite-dimensional space whose members have bounded variation and midpoint values, and let $f \geq 2B$. Define

$$\gamma_f(\mathcal{V}) = \max_{0 \neq k \in \mathcal{V}} \frac{\mathrm{Err}_f(k)}{c(\mathcal{V})\|k\|^2},$$
$$\Gamma_f(\mathcal{V}) = \max_{0 \neq k \in \mathcal{V}} \frac{\mathrm{Err}_f(k)}{\mathcal{E}_{A_f}(k)},$$
$$\Gamma_f^\eta(\mathcal{V}) = \max_{0 \neq k \in \mathcal{V}} \frac{\mathrm{Err}_f(k)}{\mathcal{E}_{A_f}(k) + \eta c(\mathcal{V})\|k\|^2}$$
$$(\eta > 0).$$

All three are finite and attained. Every $k \in \mathcal{V}$ satisfies

$$\mathrm{Err}_f(k) \leq \gamma_f(\mathcal{V}) c(\mathcal{V}) \|k\|^2,$$
$$\mathrm{Err}_f(k) \leq \Gamma_f(\mathcal{V}) \overline{\mathrm{Err}}_f(k),$$
$$\mathrm{Err}_f(k) \leq \Gamma_f^\eta(\mathcal{V}) \big(\overline{\mathrm{Err}}_f(k) + \eta c(\mathcal{V})\|k\|^2\big),$$

and $\Gamma_f^\eta(\mathcal{V})$ increases to $\Gamma_f(\mathcal{V})$ as $\eta$ decreases to 0. For two rates, $\|H_f - H_{f_0}\|_{L^2(-B,B)} \leq \left(\gamma_f^{1/2} + \gamma_{f_0}^{1/2}\right) c^{1/2} \|k\|$.

*Proof.* The error $e_f$ depends linearly on $k$, so $\mathrm{Err}_f$ and $\mathcal{E}_{A_f}$ are quadratic forms on $\mathcal{V}$, and $\mathrm{Err}_f$ is finite because $H_f$ and $K$ are bounded on $[-B,B]$. The form $\mathcal{E}_{A_f}$ is positive definite on $\mathcal{V}$ by (F2), because $A_f$ has positive measure, and $c(\mathcal{V}) > 0$.

On a finite-dimensional space, the ratio of a nonnegative quadratic form to a positive definite one attains its maximum, the largest generalized eigenvalue, and the three denominators are positive definite. The middle inequality uses $\overline{\mathrm{Err}}_f = \mathcal{E}_{A_f}$ (Theorem S4.1(c)). For each $k$ the ratio in $\Gamma_f^\eta$ increases to $\mathrm{Err}_f(k)/\mathcal{E}_{A_f}(k)$ as $\eta \downarrow 0$; exchanging the two suprema, we obtain $\lim_{\eta\to 0}\Gamma_f^\eta = \Gamma_f$. The two-rate bound is the triangle inequality applied to $e_f - e_{f_0}$. □

**Supplementary Table S4.** Fixed-grid constants of PKC spaces. The first five rows give $\gamma_f$ at the eight rates for five PKC spaces, together with its supremum over 100–500 Hz, and an entry "0" means a value below 0.005. The last three rows give $\Gamma_f^\eta$ for the main space.

| Space | $c$ | 100 | 125 | 128 | 160 | 200 | 250 | 256 | 500 Hz | sup over 100–500 Hz |
|---|---|---|---|---|---|---|---|---|---|---|
| $\mathcal{V}_{25}(D,55)$ (main) | 9.38e−3 | 0.96 | 0.05 | 0.37 | 0.13 | 0.05 | 0.04 | 0.02 | 0 | 3.72 |
| $\mathcal{V}_{22}(D,45)$ | 6.83e−2 | 0.21 | 0.07 | 0.13 | 0.05 | 0.02 | 0.08 | 0.01 | 0 | 3.09 |
| $\mathcal{V}_{25}(D,50)$ | 0.146 | 0.23 | 0.09 | 0.09 | 0.03 | 0.01 | 0.07 | 0 | 0 | 2.52 |
| $\mathcal{V}_{28}(D,55)$ | 0.500 | 0.55 | 0.06 | 0.04 | 0.01 | 0 | 0.03 | 0 | 0 | 1.07 |
| $\mathcal{V}_{31}(D,45)$ | 0.997 | 0.78 | 0.34 | 2.35 | 1.19 | 0.54 | 0.08 | 0.21 | 0 | 13.2 |
| $\Gamma_f^\eta$, main, $\eta$=$10^{-4}$ | | 1.04 | 1.41 | 3.56 | 3.59 | 3.00 | 0.82 | 1.91 | 0.06 | |
| $\Gamma_f^\eta$, main, $\eta$=$10^{-6}$ | | 1.18 | 1.51 | 4.07 | 4.23 | 4.43 | 0.89 | 4.57 | 0.95 | |
| $\Gamma_f^\eta$, main, $\eta$=$10^{-8}$ | | 1.45 | 1.54 | 5.77 | 4.67 | 4.58 | 1.33 | 4.69 | 1.21 | |

At the eight rates of Supplementary Table S4 the main space has $\gamma_f \le 0.963$, so the error of the implemented grid stays below the certificate there. Between these rates, however, the constant jumps whenever the tap set changes. Just below each rate with $Tf \in \mathbb{Z}$ the end lags $\pm T$ have left the tap set, and the response lacks the end-point term of Remark S3.1(i). For the last basis function, which has $|\varphi_{25}(\pm T; D, 55)| = 1.35$, the missing term is large: as $f \uparrow 104$ Hz, this function alone gives $\mathrm{Err}_f = 3.63c$. We computed $\gamma_f$ on a 0.01-Hz grid over 100–500 Hz, together with both one-sided limits at every rate with $Tf \in \mathbb{Z}$. For the main space the supremum is 3.72, approached as $f \uparrow 104$ Hz, compared with 1.06 at 104 Hz itself and 0.07 just above it. Every kernel of the main space therefore satisfies $\mathrm{Err}_f(k) \le 3.72c\|k\|^2$ at every rate $f \in [100,500]$ Hz. The constant exceeds 1 on $(102.986,104]$ Hz and on six shorter intervals just below 112, 120, …, 152 Hz, a total length of 3.8 Hz. The space $\mathcal{V}_{31}(D,45)$, which keeps nine functions beyond its count $N_{1/2}(D,45) = 22$, reaches 13.2 from the other side, as $f \downarrow 104$

Hz, where its end lags enter with full weight. Because the offset average is free of these jumps, the rate-uniform statements come from Theorem S4.1, whereas the implemented-grid bound of Theorem S9.1(e) uses $\gamma_f$ at the actual rate.

With $\eta = 10^{-6}$, every unit kernel of the main space satisfies $\mathrm{Err}_f(k) \leq 4.57\overline{\mathrm{Err}}_f(k) + 4.3 \times 10^{-8}$ at all eight rates. The kernels that attain $\Gamma_f^{10^{-6}}$ have end values $|k(\pm T)|$ between $3.3 \times 10^{-3}$ and $3.7 \times 10^{-2}$. Their fixed-grid errors stay below $5.1 \times 10^{-6}$, less than 1/1800 of the certificate, so the largest ratios belong to kernels whose errors are negligible. The constant $\Gamma_f$ is the limit of $\Gamma_f^{\eta}$ as $\eta \to 0$. Between $\eta = 10^{-6}$ and $10^{-8}$ the value still rises, for example from 4.07 to 5.77 at 128 Hz, while the Gram matrices are accurate to about $10^{-11}$. We therefore state the fixed-grid bound with $\Gamma_f^{\eta}$ and $\eta \geq 10^{-8}$.

(c) **Kernels that vanish at the support ends.** Let the zero extension of $k$ be absolutely continuous on $\mathbb{R}$ with $k' \in L^2(\mathbb{R})$. For $k \in H^1(I)$ this holds exactly when $k(\pm T) = 0$. Then, for $f \geq 2B$,

$$\begin{aligned}
\mathrm{Err}_f(k) &\leq \frac{\pi^2}{3}\int_{A_f} \left(\frac{|\xi| + B}{f}\right)^2 |K(\xi)|^2 d\xi \\
&\leq \frac{1}{12(f-B)^2}\int_{A_f} |\widehat{k'}(\xi)|^2 d\xi \\
&\leq \frac{\|k'\|_2^2}{12(f-B)^2}.
\end{aligned}$$

*Proof.* We have $\widehat{k'}(\xi) = 2\pi i \xi K(\xi)$. On $A_f$, $|\xi| \geq f - B$ implies $(|\xi| + B)/f \leq |\xi|/(f-B)$, which gives the second inequality, and Plancherel's theorem gives the third, so the middle integral is finite. For $|\nu| \leq B$ and $m \neq 0$, the point $\xi = \nu + mf$ satisfies $|m| \leq (|\xi| + B)/f$. The integral $\int_{-B}^{B} \sum_{m \neq 0} m^2 \, |K(\nu + mf)|^2 d\nu$ is therefore bounded by the first integral without the factor $\pi^2/3$, and the inner sum is finite for almost every $\nu$. At such $\nu$ the Cauchy–Schwarz inequality gives

$$\begin{aligned}
&\Big(\sum_{m \neq 0} |K(\nu + mf)|\Big)^2 \\
&\leq \left(\sum_{m \neq 0} m^{-2}\right) \sum_{m \neq 0} m^2 \, |K(\nu + mf)|^2 \\
&= \frac{\pi^2}{3} \sum_{m \neq 0} m^2 \, |K(\nu + mf)|^2.
\end{aligned}$$

The series of Theorem S3.1 thus converges absolutely, and $|e_f(\nu)|^2$ is bounded by the right-hand side. Integrating over $[-B, B]$, where the bands are disjoint for $f \geq 2B$, we obtain the first inequality. □

*Prior work.* The end values play the same role in the error analysis of discrete approximations of the Fourier transform, which also rests on the Poisson summation formula. When the end values of the sampled function do not match, that error is only of first order in the step (Briggs & Henson, 1995, Chapter 6; Ehler, Gröchenig & Klotz, 2026).

*Numerical check.* For three random tapered PKC kernels $u = (1 - (t/T)^2) \sum_p a_p \, \varphi_p(\cdot; D, 55)$, normalized to unit norm, the chain holds at all eight rates. At 100 Hz the first kernel gives $4.02 \times 10^{-3} \leq 1.34 \times 10^{-2} \leq 1.36 \times 10^{-2} \leq 0.958$. The bound also holds at every $\delta$ for the bumps of (a).

*Consequence for PKC.* PSWFs do not vanish at $\pm T$; the last function of the main basis has $|\varphi_{25}(\pm T; D, 55)| = 1.35$ with $\|\varphi_{25}\| = 1$. The zero extension of a PKC kernel thus jumps at $\pm T$, its derivative contains point masses there, and the right-hand side of (c) is infinite. For a random unit PKC kernel at 100 Hz, the middle integral of (c) taken over the alias bands below $\Xi$ grows in proportion to $\Xi$: it equals $2.30 \times 10^{-3}$ at $\Xi = 10^3$ Hz, $2.05 \times 10^{-2}$ at $10^4$ Hz and 0.203 at $10^5$ Hz. For PKC kernels the fixed-grid statement is therefore form (b).

### S6. Prolate kernels minimize the certificate

Proposition 2 is proved in Appendix A.3. By its part (i), PKC with $W = f_{\min} - B$ has the smallest certificate among all $P$-dimensional spaces of kernels supported in $I$. Combined with Theorem S4.1(*f*), this gives every kernel $k$ of $\mathcal{V}_P(D, f_{\min} - B)$ the spectrum-weighted bound $\sup_{f \geq f_{\min}} \overline{\mathrm{Err}}_f^{S}(k) \leq \|S\|_\infty \{1 - \lambda_P(D, f_{\min} - B)\} \|k\|^2$.

*Source.* Slepian and Pollak (1961) characterized the prolate functions by an extremal property. Among band-limited functions, the first prolate function has the largest energy concentration in a given time interval, and each later one has the largest concentration among the functions orthogonal to its predecessors. The concentration of the $p$-th function is its eigenvalue. Because the Fourier transform exchanges time and frequency, the same eigenvalues govern the concentration in $[-W, W]$ of functions supported in $I$. With $A = P_W \mathcal{F} P_I$, the two concentration operators are $A^*A$ and $AA^*$, and these share their nonzero eigenvalues. The minimax form of this property is the Courant–Fischer principle (Fischer, 1905; Courant, 1920).

*Numerical check* ($D = 0.25$ s, $W_0 = 55$ Hz). We compared the certificates of six families of $P$-dimensional spaces with the bound, and no family falls below it. The PSWF space with $W = 55$ Hz attains the bound, at $P = 15$ to within the quadrature accuracy of $2 \times 10^{-14}$.

**Supplementary Table S5.** Worst-case certificate $c(\mathcal{V})$ of six families of $P$-dimensional kernel spaces on $I$, for $D = 0.25$ s and $W_0 = 55$ Hz, next to the bound $1 - \lambda_P(D, 55)$ of Proposition 2(i).

| $P$ | $1-\lambda_P(D,55)$ | PSWF, $W$=55 | PSWF, $W$=50 | PSWF, $W$=45 | natural cubic spline, $P$ controls | Legendre, degree $<P$ | best of 100 random subspaces of $\mathcal{V}_{45}(D,55)$ |
|---|---|---|---|---|---|---|---|
| 15 | 6.3e−13 | 6.5e−13 | 2.0e−10 | 6.2e−8 | 0.217 | 0.741 | 0.781 |
| 22 | 2.48e−5 | 2.48e−5 | 1.91e−3 | 6.83e−2 | 0.344 | 0.980 | 0.904 |
| 25 | 9.38e−3 | 9.38e−3 | 0.146 | 0.511 | 0.429 | 0.996 | 0.940 |
| 28 | 0.500 | 0.500 | 0.697 | 0.893 | 0.590 | 0.9995 | 0.975 |
| 31 | 0.990 | 0.990 | 0.993 | 0.997 | 0.998 | 1.000 | 0.997 |

*Values of $P_\varepsilon(W)$ in Proposition 2(ii)* ($D = 0.25$ s, $W_0 = 55$ Hz). At every tolerance the maximum over $W$ lies at 55 Hz, and every entry is at most $N_{1-\varepsilon}(D, 55)$, which equals 25, 26, 27 and 27 for the four rows.

**Supplementary Table S6.** Largest PKC size $P_\varepsilon(W)$ whose worst-case certificate stays at or below $\varepsilon$, for $D = 0.25$ s and bands from 35 to 75 Hz.

| $\varepsilon$ | 35 | 40 | 45 | 50 | 55 | 60 | 65 | 70 | 75 Hz |
|---|---|---|---|---|---|---|---|---|---|
| 0.01 | 15 | 18 | 20 | 22 | **25** | 21 | 19 | 17 | 15 |
| 0.05 | 17 | 19 | 21 | 24 | **26** | 23 | 20 | 18 | 17 |
| 0.2 | 18 | 20 | 23 | 25 | **27** | 24 | 22 | 20 | 19 |
| 0.5 | 20 | 22 | 24 | 26 | **27** | 26 | 24 | 22 | 21 |

Proposition 2(ii) is a worst-case statement about the largest out-of-band fraction over the whole space. The error of an individual kernel depends on its own spectrum through the symbol form of Theorem S4.1(c), and Corollary S4.1(b) bounds this error when the energy beyond $W_0$ lies just above $W_0$.

**Remark S6.1 (guard band).** For $W < W_0$ the basis functions beyond the count $2DW$ place their out-of-band energy first in the guard band $\{W < |\xi| < W_0\}$, which aliases at no admissible rate. For $W = 45$ Hz, $\varphi_{23}(\cdot; D, 45)$ has 50.0% of its energy outside $[-45,45]$ Hz. The guard band 45–55 Hz holds 33.8% of the energy, and only 16.2% lies beyond 55 Hz. The certificate of a narrow band therefore crosses a given tolerance only at or after its own count. This explains why the onset for $W < W_0$ in Section 5.5 lies just above $2DW$.

### S7. The capacity-invariance limit

Lemma A.1 (coverage) and Proposition 3 are proved in Appendix A.4.

With $f_{\min} = 100$ Hz and $B = 45$ Hz, the coverage condition $f_{\max} \geq 2f_{\min} - 2B$ of Lemma A.1 reads $f_{\max} \geq 110$ Hz. The deployment range 100–500 Hz therefore covers $|\xi| \geq 55$ Hz, whereas the range 100–105 Hz leaves $(150,155)$ Hz uncovered. The eight grids used in our experiments cover $[55,4045]$ Hz, and a unit kernel of the main PKC space carries at most $4.65 \times 10^{-5}$ of its energy above 4045 Hz. Bandpass sampling theory addresses the complementary question of which rates keep the replicas of a band from overlapping (Vaughan et al., 1991).

By Proposition 3, $N_{1-\varepsilon}(D, f_{\min} - B)$ is the largest dimension of a space of kernels supported in $I$ in which no kernel places more than a fraction $\varepsilon$ of its energy in the alias region of a covering rate set. The limit depends only on $D$, $f_{\min}$ and $B$, and it holds for every linear parameterization of kernels supported in $I$, whether splines, polynomials, PSWFs or any other basis.

*Source.* Under coverage, (S1) and Lemma A.1 give $\Omega_F = \{|\xi| \geq W_0\}$, and Proposition 3 becomes the counting form of the max–min principle (Fischer, 1905; Courant, 1920) for the concentration operator $Q_{W_0}$ of Slepian and Pollak (1961). In that form, $N_{1-\varepsilon}(D, W_0)$ is the largest dimension of a subspace on which $\langle Q_{W_0}k, k\rangle \geq (1-\varepsilon)\|k\|^2$. The new step is Lemma A.1, which identifies the alias region of a covering rate set with $\{|\xi| \geq f_{\min} - B\}$ and so turns the classical count into a capacity limit for rate transfer.

*Counting the limit (classical).* Landau (1993) proved that, for $2DW \geq 2$, the eigenvalues satisfy $\lambda_{\lfloor 2DW \rfloor}(D,W) \geq \frac{1}{2} \geq \lambda_{\lfloor 2DW \rfloor + 2}(D,W)$. We use the statement as restated by Israel and Mayeli (2024, Section 1), whose indices start at 0. Hence $\lfloor 2DW \rfloor \leq \#\{p: \lambda_p \geq \frac{1}{2}\}$ and $\#\{p: \lambda_p > \frac{1}{2}\} \leq \lfloor 2DW \rfloor + 1$. When no eigenvalue equals $\frac{1}{2}$, which holds at all parameters used here, $N_{1/2}(D,W) \in \{\lfloor 2DW \rfloor, \lfloor 2DW \rfloor + 1\}$. Landau and Widom (1980) proved that, for fixed $\alpha \in (0,1)$,

$$N_\alpha(D,W) = 2DW + \frac{1}{\pi^2}\log\frac{1-\alpha}{\alpha}\log(DW) + o(\log DW) \quad (DW \to \infty),$$

where a constant factor inside the second logarithm is absorbed in the error term. Landau and Widom give no explicit bound on the remainder (Israel & Mayeli,

2024, Section 1). The formula therefore fixes the trend in $DW$, and we compute the counts used below. The dimension $2DW$ itself goes back to Landau and Pollak (1962). For $D = 0.25$ s and $W_0 = 55$ Hz, we obtain $2DW_0 = 27.5$, $N_{1/2} = 27$ and $N_{0.99} = 25$. Our computed eigenvalues satisfy Landau's bound at every $W$ from 4 to 120 Hz in steps of 0.25 Hz, which is the range where $2DW \geq 2$.

**Supplementary Table S7.** Eigenvalue counts of the time- and band-limiting operator for $D = 0.25$ s.

| $W$(Hz) | $2DW$ | $N_{0.99}$ | $N_{1/2}$ | $N_{0.05}$ | $N_{0.01}$ |
|---|---|---|---|---|---|
| 45 | 22.5 | 20 | 22 | 24 | 25 |
| 55 | 27.5 | 25 | 27 | 29 | 30 |
| 65 | 32.5 | 29 | 32 | 35 | 36 |
| 75 | 37.5 | 34 | 37 | 40 | 41 |

**Remark S7.1 (finitely many rates).** Let $F_0 = \{f_1, \ldots, f_r\}$ be rates with $f_i \geq f_{\min}$ and $\cup_i A_{f_i} \supseteq \{W_0 \leq |\xi| \leq \Xi\}$. Since $f_i \geq f_{\min} > 2B$, Theorem S4.1(c) gives $\mathcal{E}_{A_{f_i}} = \overline{\mathrm{Err}}_{f_i}$, so $\mathcal{E}_{[W_0,\Xi]} \leq \sum_i \overline{\mathrm{Err}}_{f_i}$. Hence every $k$ satisfies $\max_i \overline{\mathrm{Err}}_{f_i}(k) \geq \frac{1}{r}\{\mathcal{E}_{\geq W_0}(k) - \mathcal{E}_{>\Xi}(k)\}$. The max–min step of Proposition 2(i) needs no coverage, so a space with $P > N_{1-\varepsilon}(D, W_0)$ contains a kernel whose offset-averaged error at one of the $r$ grids exceeds $\{\varepsilon\|k\|^2 - \mathcal{E}_{>\Xi}(k)\}/r$.

#### S8. Full-rank realization

**Theorem S8.1 (realization rank) (classical).** Let $\ell_f = 2\lfloor Tf \rfloor + 1$ be the number of lags in $I$ at rate $f$, and let $M_f \in \mathbb{R}^{\ell_f \times P}$ have the entries $(M_f)_{np} = w_n \varphi_p(n/f; D, W)$, so that the taps of $k = \sum_p a_p \varphi_p$ are $M_f a/f$.

(a) $\mathrm{rank} M_f = \min\{P, \ell_f\}$. For $P \leq \ell_f$ the matrix has full column rank, and for $P > \ell_f$ its null space has dimension $P - \ell_f$.

(b) If $P \leq \ell_{f_0}$, the training map $a \mapsto \sum_p a_p H_{f_0}[\varphi_p]$, restricted to any set of frequencies of positive measure, is injective. In particular, no nonzero coefficient vector leaves the realized response on the data band unchanged at the training rate.

*Source.* For every $N$, the first $N$ prolate functions form a Chebyshev system on the closed interval (Hogan & Lakey, 2012, Theorem 2.1.16 and Lemma 2.1.2, as used by Hogan & Lakey, 2015, Section 2). Thus $\det[\varphi_p(t_i)]_{i,p \leq N} \neq 0$ for any points $t_1 < \cdots < t_N$ in $I$.

*Proof.* (a) If $P \leq \ell_f$, we choose $P$ of the $\ell_f$ lags. By the Chebyshev property with $N = P$, the corresponding $P \times P$ matrix $[\varphi_p(t_i)]$ is nonsingular, and multiplying its rows by the positive weights $w_n \in \{\frac{1}{2}, 1\}$ keeps it nonsingular. If $P > \ell_f$, the Chebyshev property with $N = \ell_f$ makes the submatrix formed by the first $\ell_f$ columns nonsingular, so the rank is $\ell_f$ and the null space has dimension $P - \ell_f$. (b) The response $\nu \mapsto \sum_n h_n \, e^{-2\pi i \nu n/f_0}$ is a trigonometric polynomial and extends to an entire function. If it vanishes on a set of positive measure, it vanishes identically, so its taps $h = M_{f_0} a/f_0$ vanish, and $a = 0$ by (a). □

*Numerical check.* We evaluated $\det[\varphi_p(t_i)]$ in 60-digit arithmetic over 300 random ordered point sets for each of $(W, P) = (55{,}5)$, $(55{,}25)$ and $(45{,}31)$ in Hz. Half of the sets contained both end points, and the determinant kept a single sign in every case. Extended precision is needed here because the determinants reach $10^{-54}$ and the end values of low-order PSWFs fall below $10^{-11}$. Over nine PKC spaces and eight rates, the realization matrix has full column rank whenever $P \le \ell_f$, and otherwise its null space has dimension $P - \ell_f$, for example 6 for $(45{,}31)$ at 100 Hz. For the main space, $M_f/\sqrt{f}$ has smallest singular value 0.240 at 100 Hz, where $\ell_f = P = 25$, and at least 0.991 at the other seven rates. The condition number separates the spaces further. It stays at most 5.9 for the main space at all eight rates, whereas the space with $W = 45$ Hz and $P = 31$ reaches $4.1 \times 10^3$ at 125 Hz, where $\ell_f = P = 31$. On $[-45{,}45]$ Hz the training map of the main space has largest singular value 1.00, and its smallest singular value is 0.060 at 200 Hz and 0.095 at 100 Hz. The smallest singular value measures how strongly the data band determines the weakest coefficient direction.

### S9. Cascades of continuous-kernel layers

**Theorem S9.1 (cascade bound) (new).** Consider $L$ linear time-invariant layers with matrix kernels $k_l : \mathbb{R} \to \mathbb{C}^{C_l \times C_{l-1}}$ whose entries have bounded variation, midpoint values and support in $[-T_l, T_l]$, and whose entries lie in a finite-dimensional space $\mathcal{V}_l$. Every layer is realized entrywise at the same rate $f$, with an offset $\theta_l$ shared by its entries ($\theta_l = 0$ in the implementation), and $H_{f,\theta_l,l}$ denotes its response referred to physical time. Assume A1 for the network input. Let $\alpha_l = \sup_{|\nu| \le B} \|K_l(\nu)\|_2$ and $\beta_l(f) = \sup_{\theta, |\nu| \le B} \|H_{f,\theta,l}(\nu)\|_2$ (spectral norms), and for matrix-valued $U$ let $\|U\|_S^2 = \int_{-B}^{B} S \, \|U\|_F^2$. Write $\Pi_{f,\theta} = H_{f,\theta_L,L} \cdots H_{f,\theta_1,1}$, $\Pi_f = \Pi_{f,0}$ and $\Pi = K_L \cdots K_1$, let $e_{f,\theta_l,l} = H_{f,\theta_l,l} - K_l$, and let

$$\chi_l(f) = \prod_{i<l} \alpha_i \prod_{i>l} \beta_i(f),$$

$$\chi_l = \prod_{i<l} \alpha_i \prod_{i>l} \max\{\beta_i(f), \beta_i(f_0)\}.$$

Energies and errors of matrix kernels are summed over the entries: $\mathcal{E}_\Omega(k_l) = \sum_{a,b} \mathcal{E}_\Omega((k_l)_{ab})$, $\overline{\mathrm{Err}}_f^S(k_l) = \sum_{a,b} \overline{\mathrm{Err}}_f^S((k_l)_{ab})$ and $\|k_l\|_F^2 = \sum_{a,b} \|(k_l)_{ab}\|^2$.

(a) The realized cascade maps an input with spectrum $X$ to the samples of the band-limited signal with spectrum $e^{2\pi i\nu \sum_l \theta_l/f}\Pi_{f,\theta}X$. This signal is $\Pi_{f,\theta}X$ shifted in time by the known amount $\sum_l \theta_l/f$, whereas the continuous cascade gives $\Pi X$. Parts (b)–(*f*) compare the shift-compensated output $\Pi_{f,\theta}X$ with $\Pi X$, and the implemented offsets $\theta_l = 0$ produce no shift.

(b) (One rate, fixed offsets.) $\|\Pi_{f,\theta} - \Pi\|_S \le \sum_{l=1}^L \chi_l(f)\|e_{f,\theta_l,l}\|_S$.

(c) (One rate, random offsets.) Let each $\theta_l$ be uniform on $[0,1)$, with any dependence between layers. Then

$$(\mathbb{E}\|\Pi_{f,\theta} - \Pi\|_S^2)^{1/2} \le \sum_l \chi_l(f)\overline{\mathrm{Err}}_f^S(k_l)^{1/2}$$
$$\le \|S\|_\infty^{1/2} \sum_l \chi_l(f)\mathcal{E}_{A_f}(k_l)^{1/2}.$$

(d) (Two rates.) For $f, f_0 \ge f_{\min}$ and offsets $\theta$ at $f$ and $\theta'$ at $f_0$ as in (c),

$$(\mathbb{E}\|\Pi_{f,\theta} - \Pi_{f_0,\theta'}\|_S^2)^{1/2} \le \|S\|_\infty^{1/2} \sum_l \chi_l \left(\mathcal{E}_{A_f}(k_l)^{1/2} + \mathcal{E}_{A_{f_0}}(k_l)^{1/2}\right)$$
$$\le 2\|S\|_\infty^{1/2} \sum_l \chi_l\, c(\mathcal{V}_l)^{1/2}\|k_l\|_F.$$

(e) (Implemented grids.)

$$\|\Pi_f - \Pi_{f_0}\|_S \le \sum_l \chi_l \left(\|e_{f,0,l}\|_S + \|e_{f_0,0,l}\|_S\right)$$
$$\le \|S\|_\infty^{1/2} \sum_l \chi_l \left(\gamma_f(\mathcal{V}_l)^{1/2} + \gamma_{f_0}(\mathcal{V}_l)^{1/2}\right)c(\mathcal{V}_l)^{1/2}\|k_l\|_F.$$

(*f*) If every layer is non-expansive on the data band ($\alpha_l \le 1$ and $\beta_l \le 1$), the root-mean-square rate-transfer error of the cascade over random offsets is at most $2\|S\|_\infty^{1/2} \sum_l c(\mathcal{V}_l)^{1/2}\|k_l\|_F$, which is twice the sum of the per-layer certificates.

*Proof.* (a) A layer realized with offset $\theta_l$ is the discrete filter with taps $h_{f,\theta_l}[n]$ at integer sample lags, whose transfer function is $e^{2\pi i\nu\theta_l/f}H_{f,\theta_l,l}(\nu)$. By Lemma S2.1 it maps the samples of a signal band-limited to $[-B, B]$ to the samples of the signal whose spectrum is multiplied by this transfer function, which is again band-limited to $[-B, B]$. Induction over $l$ gives (a), and removing the known shift $\sum_l \theta_l / f$ from the realized output leaves $\Pi_{f,\theta} X$. (b) We start from the telescoping identity

$$\Pi_{f,\theta} - \Pi = \sum_{l=1}^{L} H_{f,\theta_L,L} \cdots H_{f,\theta_{l+1},l+1} \big(H_{f,\theta_l,l} - K_l\big) K_{l-1} \cdots K_1.$$

The pointwise inequality $\|UeV\|_F \le \|U\|_2 \|e\|_F \|V\|_2$ bounds the norm of the $l$-th term by $\chi_l(f)\|e_{f,\theta_l,l}\|_S$, and the triangle inequality gives (b). (c) Minkowski's inequality in $L^2(\mathbb{P} \otimes S dv)$, where $\mathbb{P}$ is the law of the offsets, bounds the root-mean-square norm of the sum by the sum of the root-mean-square norms of the terms. The $l$-th term depends on all offsets, but its bound $\chi_l(f)\|e_{f,\theta_l,l}\|_S$ depends on $\theta_l$ only, because $\beta_i$ is a supremum over offsets. Since $\theta_l$ is uniform and the offset is shared within the layer, Theorem S4.1(c), applied entrywise and summed, gives $\mathbb{E}\|e_{f,\theta_l,l}\|_S^2 = \overline{\mathrm{Err}}_f^S(k_l)$, and Theorem S4.1(*f*) bounds it by $\|S\|_\infty \mathcal{E}_{A_f}(k_l)$. (d) We write $\Pi_{f,\theta} - \Pi_{f_0,\theta'} = (\Pi_{f,\theta} - \Pi) - (\Pi_{f_0,\theta'} - \Pi)$, apply Minkowski's inequality and (c) at both rates with $\chi_l(f), \chi_l(f_0) \le \chi_l$, and use $\mathcal{E}_{A_f}(k_l) \le \mathcal{E}_{\ge W_0}(k_l) \le c(\mathcal{V}_l)\|k_l\|_F^2$ from (S1). (e) We apply (b) at $f$ and at $f_0$ with zero offsets, use the triangle inequality through $\Pi$, and bound each entry by Proposition S5.1(b). (*f*) follows from (d). □

*Scope.* Theorem S9.1 covers linear stacks whose layers all run at one rate. A channel-mixing layer such as the spatial projection has a lag-0 matrix kernel, which is a point mass and falls outside the bounded-variation hypothesis. We therefore treat it as an exact layer, applied with no offset and no shift. Its response is the constant matrix $K_l = H_{f,0,l}$, so $e_{f,0,l} = 0$, its term drops out of (b)–(e), and it enters only through $\alpha_l = \beta_l = \|K_l\|_2$. In the factorized backbone of our experiments, the spatial projection and the temporal layer form the linear part, and the log-variance head acts after it. The theorem with $L = 2$ thus bounds the rate-transfer change of every map that enters the head. In a deeper stack with pointwise nonlinearities between temporal layers, the theorem applies to a linear segment only when its input satisfies A1. A pointwise nonlinearity widens the band and therefore generally violates this assumption.

*Prior work.* The telescoping identity for a product of perturbed operators is standard. For Fourier neural operators, Lanthaler, Stuart and Trautner (2024) bound the error of the grid implementation in terms of the grid resolution and the regularity of the input. Theorem S9.1 instead bounds the rate-transfer error of a cascade by per-layer certificates computed from the weights.

*Numerical check.* We stacked three layers with $2 \times 2$ matrix kernels whose entries were random members of the main PKC space, scaled each layer to $\alpha_l = 1$, and compared five deployment rates with $f_0 = 200$ Hz. Root-mean-square values were taken over 300 independent draws of the offsets, and the estimated $\beta_l$ were at most 1.008. All bounds hold at the five rates, as Supplementary Table S8 shows.

**Supplementary Table S8.** Root-mean-square errors of the three-layer cascade and the bounds of Theorem S9.1 at five deployment rates.

| $f$(Hz) | RMS $\|\Pi_{f,\theta} - \Pi\|_S$ | bound (c) | RMS $\|\Pi_{f,\theta} - \Pi_{200,\theta'}\|_S$ | first bound of (d) | $\|\Pi_f - \Pi_{200}\|_S$ | first bound of (e) |
|---|---|---|---|---|---|---|
| 100 | 0.101 | 0.512 | 0.104 | 0.654 | 0.116 | 0.625 |
| 125 | 0.045 | 0.255 | 0.050 | 0.398 | 0.041 | 0.234 |
| 160 | 0.032 | 0.184 | 0.040 | 0.326 | 0.012 | 0.306 |
| 250 | 0.019 | 0.112 | 0.032 | 0.255 | 0.036 | 0.217 |
| 500 | 0.009 | 0.055 | 0.027 | 0.197 | 0.021 | 0.126 |

The certificate forms, that is, the second bounds of (d) and (e), range from 4.29 to 4.30 and from 0.52 to 2.60, respectively. They exceed the first bounds because random kernels of the main space place far less than the maximal fraction $c$ of their energy beyond 55 Hz. For an isotropic random coefficient vector, the expected fraction is $\frac{1}{25}\sum_{p\leq 25}(1 - \lambda_p(D, 55))$, about $c/21$.

### S10. General kernels under two realization rules

In Table 2, the general continuous kernels are sampled at the exact lags $n/f$ with trapezoid end weights, which is the rule of ProlateNet. Their own layers instead place the taps evenly over the support (linspace taps), and this rescales time whenever $Df$ is not an even integer (Section 4.4). We trained and evaluated each general kernel under both rules with the protocol of Section 5, and Supplementary Table S9 lists the results. The exact-lag rule lowers the mean worst drop of Spline-15, Spline-31 and Sinc on both EEG datasets, and Spline-15 is stable on both only under this rule.

**Supplementary Table S9.** Score at the training rate / mean worst drop over all target rates for the general continuous kernels under two realization rules. Bold cells are stable and accurate in the sense of Section 5.2.

| Design, realization | SEED, macro-F1 | Sleep-EDF, macro-F1 | PTB-XL, 100 Hz, macro-AUROC | PTB-XL, 500 Hz, macro-AUROC |
|---|---|---|---|---|
| Spline-15, exact lags with trapezoid end weights | **0.544 / 0.0053** | **0.713 / 0.0036** | **0.893 / 0.0004** | **0.891 / 0.0007** |
| Spline-15, linspace taps | 0.540 / 0.0170 | 0.714 / 0.0233 | **0.892 / 0.0032** | **0.893 / 0.0042** |

| Design, realization | SEED, macro-F1 | Sleep-EDF, macro-F1 | PTB-XL, 100 Hz, macro-AUROC | PTB-XL, 500 Hz, macro-AUROC |
|---|---|---|---|---|
| Spline-31, exact lags with trapezoid end weights | 0.528 / 0.0120 | **0.710 / 0.0039** | **0.891 / 0.0004** | **0.893 / 0.0016** |
| Spline-31, linspace taps | 0.540 / 0.0253 | 0.712 / 0.0179 | **0.893 / 0.0087** | 0.893 / 0.0139 |
| Spline-63, exact lags with trapezoid end weights | 0.542 / 0.1341 | 0.712 / 0.5399 | 0.892 / 0.1130 | 0.893 / 0.0780 |
| Spline-63, linspace taps | 0.535 / 0.1525 | 0.711 / 0.5004 | 0.892 / 0.1114 | 0.892 / 0.0681 |
| Sinc, exact lags with trapezoid end weights | **0.525 / 0.0006** | 0.689 / 0.0002 | 0.882 / −0.0001 | 0.883 / 0.0001 |
| Sinc, linspace taps | **0.525 / 0.0077** | 0.685 / 0.0040 | 0.882 / 0.0005 | 0.884 / 0.0005 |
| SIREN, exact lags with trapezoid end weights | 0.494 / 0.1356 | 0.582 / 0.1154 | 0.856 / 0.0198 | 0.856 / 0.0191 |
| SIREN, linspace taps | 0.502 / 0.1312 | 0.595 / 0.1390 | 0.855 / 0.0174 | 0.858 / 0.0250 |

### S11. Statistical tests for Table 2

A design is stable when its mean worst drop stays below 0.01. It is accurate when its training-rate score is not significantly below that of the CNN in a paired two-sided test at $\alpha = 0.05$. On SEED and Sleep-EDF, the test is a Wilcoxon signed-rank test over fold means. PTB-XL has a single official test fold, so its test is a paired bootstrap over the 2,158 test records with 2,000 resamples, in which the same records enter every design and each design's score is averaged over its five seeds. Supplementary Table S10 lists both tests for every design and column of Table 2. ProlateNet never scores significantly below the CNN at the training rate. Scores significantly above that of the CNN occur only on PTB-XL, among them ProlateNet at 500 Hz. Drops significantly smaller than that of ProlateNet occur for Sinc on SEED and on PTB-XL at 100 Hz and for the augmented CNN on PTB-XL at 500 Hz.

**Supplementary Table S10.** p values of the paired two-sided tests behind Table 2, given as score / drop. Score: training-rate score against the CNN; drop: mean worst drop against ProlateNet. SEED and Sleep-EDF: Wilcoxon signed-rank tests on fold means; PTB-XL: paired bootstrap over the 2,158 test records with 2,000 resamples, where the smallest attainable p is 0.001. Bold: $p < 0.05$.

| Design | SEED | Sleep-EDF | PTB-XL, 100 Hz | PTB-XL, 500 Hz |
|---|---|---|---|---|
| ProlateNet | 0.12 / — | 0.92 / — | 0.30 / — | **0.011** / — |
| CNN | — / **6.1 × 10$^{-5}$** | — / **0.0020** | — / **0.001** | — / **0.001** |
| Spline-15 | 0.64 / **0.0015** | 0.625 / 0.23 | **0.029** / 0.10 | **0.048** / **0.0050** |
| Spline-31 | 0.055 / **6.1 × 10$^{-5}$** | 0.19 / 0.625 | 0.49 / 0.24 | **0.001** / **0.001** |
| Spline-63 | 1 / **6.1 × 10$^{-5}$** | 0.85 / **0.0020** | 0.11 / **0.001** | **0.001** / **0.001** |
| Sinc | 0.073 / **6.1 × 10$^{-5}$** | **0.0020** / 0.16 | **0.001** / **0.021** | **0.0020** / 0.64 |
| SIREN | **0.0026** / **6.1 × 10$^{-5}$** | **0.0020** / **0.0039** | **0.001** / **0.001** | **0.001** / **0.001** |
| Cosine basis | 0.23 / **6.1 × 10$^{-5}$** | 0.85 / **0.0039** | 0.10 / **0.001** | **0.0050** / **0.001** |
| FlexConv | 0.055 / **6.1 × 10$^{-5}$** | **0.0059** / **0.049** | **0.0080** / **0.001** | 0.66 / **0.001** |
| FlexConv-fit | **0.026** / 0.083 | **0.014** / **0.0098** | **0.001** / **0.001** | **0.001** / **0.001** |
| FlexConv-fit (55 Hz) | 0.064 / **1.2 × 10$^{-4}$** | **0.0039** / **0.0020** | **0.001** / **0.001** | **0.001** / **0.001** |
| SFI | 0.45 / **6.1 × 10$^{-5}$** | 0.32 / **0.0020** | **0.001** / **0.001** | **0.001** / **0.001** |

| Design | SEED | Sleep-EDF | PTB-XL, 100 Hz | PTB-XL, 500 Hz |
|---|---|---|---|---|
| SFI-band | **0.026** / **0.048** | 0.56 / **0.0039** | **0.001** / **0.001** | **0.001** / **0.001** |
| SFI-band (55 Hz) | 0.17 / **$6.1 \times 10^{-5}$** | 0.32 / **0.0020** | **0.001** / **0.001** | **0.001** / **0.001** |
| *Multi-rate augmentation* | | | | |
| CNN | **0.022** / **$6.1 \times 10^{-5}$** | 0.064 / 0.084 | **0.001** / **0.001** | **0.001** / **0.025** |
| Spline-31 | **0.012** / **0.0012** | 0.625 / 0.56 | 0.11 / 0.39 | **0.001** / **0.035** |

**References cited in the Supplementary Material**


Blu, T., & Unser, M. (1999). Quantitative Fourier analysis of approximation techniques. I. Interpolators and projectors. *IEEE Transactions on Signal Processing*, 47(11), 2783–2795. https://doi.org/10.1109/78.790659

Briggs, W. L., & Henson, V. E. (1995). *The DFT: An Owner's Manual for the Discrete Fourier Transform*. Society for Industrial and Applied Mathematics. https://doi.org/10.1137/1.9781611971514

Brown, J. L., Jr. (1978). On mean-square aliasing error in the cardinal series expansion of random processes (Corresp.). *IEEE Transactions on Information Theory*, 24(2), 254–256. https://doi.org/10.1109/TIT.1978.1055846

Courant, R. (1920). Über die Eigenwerte bei den Differentialgleichungen der mathematischen Physik. *Mathematische Zeitschrift*, 7, 1–57. https://doi.org/10.1007/BF01199396

Cranley, R., & Patterson, T. N. L. (1976). Randomization of number theoretic methods for multiple integration. *SIAM Journal on Numerical Analysis*, 13(6), 904–914. https://doi.org/10.1137/0713071

Crochiere, R. E., & Rabiner, L. R. (1975). Optimum FIR digital filter implementations for decimation, interpolation, and narrow-band filtering. *IEEE Transactions on Acoustics, Speech, and Signal Processing*, 23(5), 444–456. https://doi.org/10.1109/TASSP.1975.1162719

Cruz-Orive, L. M. (1989). On the precision of systematic sampling: A review of Matheron's transitive methods. *Journal of Microscopy*, 153(3), 315–333. https://doi.org/10.1111/j.1365-2818.1989.tb01480.x

Dirichlet, P. G. L. (1829). Sur la convergence des séries trigonométriques qui servent à représenter une fonction arbitraire entre des limites données. *Journal für die reine und angewandte Mathematik*, 4, 157–169. https://doi.org/10.1515/crll.1829.4.157

Ehler, M., Gröchenig, K., & Klotz, A. (2026). Quantitative estimates: How well does the discrete Fourier transform approximate the Fourier transform on ℝ? *SIAM Review*, 68(1), 93–123. https://doi.org/10.1137/24M1650399

Fischer, E. (1905). Über quadratische Formen mit reellen Koeffizienten. *Monatshefte für Mathematik und Physik*, 16, 234–249. https://doi.org/10.1007/BF01693781

Hogan, J. A., & Lakey, J. D. (2012). *Duration and Bandwidth Limiting: Prolate Functions, Sampling, and Applications*. Birkhäuser (Applied and Numerical Harmonic Analysis). https://doi.org/10.1007/978-0-8176-8307-8

Hogan, J. A., & Lakey, J. D. (2015). Frame properties of shifts of prolate spheroidal wave functions. *Applied and Computational Harmonic Analysis*, 39(1), 21–32. https://doi.org/10.1016/j.acha.2014.08.003

Israel, A., & Mayeli, A. (2024). On the eigenvalue distribution of spatio-spectral limiting operators in higher dimensions. *Applied and Computational Harmonic Analysis*, 70, 101620. https://doi.org/10.1016/j.acha.2023.101620

Jackson, L. B. (2000). A correction to impulse invariance. *IEEE Signal Processing Letters*, 7(11), 273–275. https://doi.org/10.1109/97.870677

Jordan, C. (1881). Sur la série de Fourier. *Comptes rendus hebdomadaires des séances de l'Académie des sciences*, 92, 228–230. No DOI.

Landau, H. J. (1993). On the density of phase-space expansions. *IEEE Transactions on Information Theory*, 39(4), 1152–1156. https://doi.org/10.1109/18.243434

Landau, H. J., & Pollak, H. O. (1962). Prolate spheroidal wave functions, Fourier analysis and uncertainty, III: The dimension of the space of essentially time- and band-limited signals. *Bell System Technical Journal*, 41(4), 1295–1336. https://doi.org/10.1002/j.1538-7305.1962.tb03279.x

Landau, H. J., & Widom, H. (1980). Eigenvalue distribution of time and frequency limiting. *Journal of Mathematical Analysis and Applications*, 77(2), 469–481. https://doi.org/10.1016/0022-247X(80)90241-3

Lanthaler, S., Stuart, A. M., & Trautner, M. (2024). Discretization error of Fourier neural operators. arXiv:2405.02221. https://doi.org/10.48550/arXiv.2405.02221

L'Ecuyer, P., & Lemieux, C. (2000). Variance reduction via lattice rules. *Management Science*, 46(9), 1214–1235. https://doi.org/10.1287/mnsc.46.9.1214.12231

Matheron, G. (2019). *Matheron's Theory of Regionalised Variables* (V. Pawlowsky-Glahn & J. Serra, Eds.). Oxford University Press. Lecture notes of the 1970 Fontainebleau summer school, first issued as Fascicule 5 of *Les Cahiers du Centre de Morphologie Mathématique de Fontainebleau*; cited as Matheron (1970/2019). https://doi.org/10.1093/oso/9780198835660.001.0001

Mecklenbräuker, W. F. G. (2000). Remarks on and correction to the impulse invariant method for the design of IIR digital filters. *Signal Processing*, 80(8), 1687–1690. https://doi.org/10.1016/S0165-1684(00)00113-4

Moran, P. A. P. (1950). Numerical integration by systematic sampling. *Mathematical Proceedings of the Cambridge Philosophical Society*, 46(1), 111–115. https://doi.org/10.1017/S0305004100025524

Park, S. K., & Schowengerdt, R. A. (1982). Image sampling, reconstruction, and the effect of sample-scene phasing. *Applied Optics*, 21(17), 3142–3151. https://doi.org/10.1364/AO.21.003142

Poisson, S.-D. (1823). Suite du mémoire sur les intégrales définies et sur la sommation des séries. *Journal de l'École Royale Polytechnique*, 12 (cahier 19), 404–509. No DOI.

Poisson, S.-D. (1827). Mémoire sur le calcul numérique des intégrales définies. *Mémoires de l'Académie royale des sciences de l'Institut de France*, 6 (année 1823), 571–602. No DOI.

Shannon, C. E. (1949). Communication in the presence of noise. *Proceedings of the IRE*, 37(1), 10–21. https://doi.org/10.1109/JRPROC.1949.232969

Slepian, D., & Pollak, H. O. (1961). Prolate spheroidal wave functions, Fourier analysis and uncertainty, I. *Bell System Technical Journal*, 40(1), 43–63. https://doi.org/10.1002/j.1538-7305.1961.tb03976.x

Vaughan, R. G., Scott, N. L., & White, D. R. (1991). The theory of bandpass sampling. *IEEE Transactions on Signal Processing*, 39(9), 1973–1984. https://doi.org/10.1109/78.134430

Venkataramani, R., & Bresler, Y. (2000). Perfect reconstruction formulas and bounds on aliasing error in sub-Nyquist nonuniform sampling of multiband signals. *IEEE Transactions on Information Theory*, 46(6), 2173–2183. https://doi.org/10.1109/18.868487

Zygmund, A. (2003). *Trigonometric Series* (3rd ed., Vols. I and II, with a foreword by R. Fefferman). Cambridge University Press. https://doi.org/10.1017/CBO9781316036587
